## Supplementary material for "A stochastic RNA editing process targets a select number of sites in individual *Drosophila* glutamatergic motoneurons": MATLAB code: process_flybase_data.m

```
% A stochastic RNA editing process targets a limited number of sites in individual Drosophila glutamatergic motoneurons
% Andrés B. Crane, Suresh K. Jetti, J. Troy Littleton
% The Picower Institute for Learning and Memory, Department of Brain and Cognitive Sciences, Department of Biology, Massachusetts Institute of Technology, Cambridge, MA 02139
% Correspondence and requests for materials should be addressed to J.T.L..


%% This code creates the flybase_data file
% flybase_data contains the start and end locations of each mRNA transcript
% listed in flybase

%Download these files from http://ftp.flybase.net/genomes/Drosophila_melanogaster/dmel_r6.52_FB2023_03/fasta/
files = ["dmel-all-CDS-r6.51.fasta";...
             "dmel-all-five_prime_UTR-r6.51.fasta";...
             "dmel-all-three_prime_UTR-r6.51.fasta"];
types = ["CDS";...
         "FivePrimeUTR";...
         "ThreePrimeUTR"];

path.main = fileparts(cd);
path.genepartdata = [path.main '\Flybase genomic data\GenePart data\'];

addpath(path.genepartdata)

tic    

flybase_data = struct();
    currentFastaTypeArray = {};
%preallocated memory for cell arrays
%     transcriptIDArray = cell(fastaSize,size(types,1));
%     geneNameArray = cell(fastaSize,1);
transcriptIDArray = {};
     geneNameArray = {};

%for each of the fasta files:
for currentFasta = 1:size(files,1)
    currentFile = files(currentFasta);
    currentType = types(currentFasta);
    

    %load fasta file
    f = waitbar(0,strcat("Loading ",currentType," Fasta"));
    fasta = fastaread(currentFile);
    fastaSize = size(fasta,1);
    
    %update waitbar
    waitbar(0,f,strcat("Extracting ", currentType))

    %for each entry in the fasta:
    for i =1:fastaSize

        % Split the header into chunks
        C = strsplit(fasta(i).Header, {' ',';'});

        % Determine if gene is on + or - ("complement") strand
        index_complement = contains(C,"complement");

        if sum(index_complement)==0
            dnaStrand = "+";
        elseif sum(index_complement)>0
            dnaStrand = "-";
        end

        % Find the location box, and the next one will have the genomic
        % positions
        indeces = contains(C,"loc=");

        % Break up the positions into cells
        try
            positions = strsplit(C{indeces}, {'..', ',', '(', ')',':'});
        catch
            i
            currentFasta
            fasta(i).Header
        end
        positions = str2double(positions);
        numbersIdx = ~isnan(positions);
        positions = positions(numbersIdx);
        fasta_data = reshape(positions,[2, numel(positions)/2]);
        fasta_data = num2cell(fasta_data');

        previousEndPosition = 0;
        for DNAsegment = 1:size(fasta_data,1)
            endPosition = fasta_data{DNAsegment,2}-fasta_data{DNAsegment,1}+previousEndPosition+1;
            startPosition = previousEndPosition+1;
            fasta_data(DNAsegment,3) = extractBetween(fasta(i).Sequence,startPosition,endPosition);
            previousEndPosition = endPosition;
        end

        if currentType == "CDS"
            geneNameArray{i} = C{1};
        end

        % Accumulate data into an array
        geneID = extract(fasta(i).Header,"parent=FBgn"+digitsPattern(7));
        geneId = extractAfter(geneID,'=');
        geneIDArray{i,currentFasta} = string(geneId);

        transcriptID = extract(fasta(i).Header,"FBtr"+digitsPattern(7));
        transcriptIDArray{i,currentFasta} = string(transcriptID);
        currentFastaTypeArray{i,currentFasta} = fasta_data;
        dnaStrandArray{i,currentFasta} = dnaStrand;
        
        
        
        %Update waitbar
        if mod(i,50)==0
            waitbar(i/fastaSize,f,strcat("Extracting ", currentType))
        end

    end

    close(f)
end
    toc

%Match transcripts on the correct rows
% Figure out how to incorporate dnaStrandArray!

% Find indeces for non-empty rows
indeces2 = ~cellfun(@isempty,transcriptIDArray(:,2));

% Compare non-empty rows for first two fasta types
% Locb2 are indeces to use on IDarray col 2 to get to to match col 1
[~,Locb2] = ismember(string(transcriptIDArray(indeces2,2)),string(transcriptIDArray(:,1)));

indeces3 = ~cellfun(@isempty,transcriptIDArray(:,3));
[~,Locb3] = ismember(string(transcriptIDArray(indeces3,3)),string(transcriptIDArray(:,1)));

transcriptIDArrayMatchedRows = {};
transcriptIDArrayMatchedRows(:,1) = currentFastaTypeArray(:,1);
transcriptIDArrayMatchedRows(Locb2,2) = currentFastaTypeArray(indeces2,2);
transcriptIDArrayMatchedRows(Locb3,3) = currentFastaTypeArray(indeces3(1:size(Locb3,1)),3);

transcriptIDArrayMatchedRows(:,4) = dnaStrandArray(:,1);


flybase_data = struct("Gene_Name",geneNameArray',...
                "GeneID",geneIDArray(:,1),...
                "TranscriptID",transcriptIDArray(:,1),...
                types(1),transcriptIDArrayMatchedRows(:,1),...
                types(2),transcriptIDArrayMatchedRows(:,2),...
                types(3),transcriptIDArrayMatchedRows(:,3),...
                "DNAstrand",transcriptIDArrayMatchedRows(:,4));


% Save data
save([path.genepartdata 'flybase_data_2023-07-20_r6.51.mat'],"flybase_data")
```
