## Supplementary material for "A stochastic RNA editing process targets a select number of sites in individual *Drosophila* glutamatergic motoneurons": MATLAB code: Step1_Import.m

```
% A stochastic RNA editing process targets a limited number of sites in individual Drosophila glutamatergic motoneurons
% Andrés B. Crane, Suresh K. Jetti, J. Troy Littleton
% The Picower Institute for Learning and Memory, Department of Brain and Cognitive Sciences, Department of Biology, Massachusetts Institute of Technology, Cambridge, MA 02139
% Correspondence and requests for materials should be addressed to J.T.L..


%% Set up working directory and MATLAB environment
clear
clc
path.main = 'C:\Users\Littleton Lab\Documents\Andres C drive\RNA editing\GATK analysis - Muscle vs neurons\';
path.m = [path.main '\GATK output\Muscle vs Neuron\'];


%% Import data
% Set up the Import Options and import the data
opts = delimitedTextImportOptions("NumVariables", 13);

% Specify range and delimiter
opts.DataLines = [2, Inf];
opts.Delimiter = "\t";

% Specify column names and types
opts.VariableNames = ["CellID", "Type", "Edit", "CHROM", "POS", "ID", "REF", "ALT", "QUAL", "FILTER", "INFO", "FORMAT", "DETAILS"];
opts.VariableTypes = ["categorical", "string", "double", "string", "double", "categorical", "categorical", "categorical", "double", "categorical", "string", "string", "string"];

% Specify file level properties
opts.ExtraColumnsRule = "ignore";
opts.EmptyLineRule = "read";

% Specify variable properties
opts = setvaropts(opts, ["INFO", "DETAILS"], "WhitespaceRule", "preserve");
opts = setvaropts(opts, ["CellID", "Type", "CHROM", "ID", "REF", "ALT", "FILTER", "INFO", "FORMAT", "DETAILS"], "EmptyFieldRule", "auto");

% Import the data
Ib = readtable([path.m 'Ib.filtered'], opts);
Is = readtable([path.m 'Is.filtered'], opts);
M = readtable([path.m 'M.filtered'], opts);
masterTable = [Ib;Is;M];


% Clear temporary variables
clear opts

%% Clean up imported data

% Delete final row with undefined data
%masterTable(end,:) = [];

% Get rid of <non-ref> allele in ALT column
altcol = string(masterTable.ALT);
newalt = split(altcol,",");
masterTable.ALT = categorical(newalt(:,1));

% Remove ID column
masterTable.ID = [];

% Remove FILTER column
masterTable.FILTER = [];

% Create DP column
infocol = masterTable.INFO;
newpat = lookBehindBoundary("DP=");
dpinfo = extract(infocol,newpat+digitsPattern);
masterTable.Depth = double(dpinfo(:,1));
masterTable = movevars(masterTable,'Depth','After','ALT');

% Combine Chr and pos into one variable
chrcol = masterTable.CHROM;
poscol = masterTable.POS;
masterTable.('Edit Position') = append(string(chrcol),":",string(poscol));
masterTable = movevars(masterTable,'Edit Position','After','Type');

% Rename Type to CellType
masterTable = renamevars(masterTable,"Type","CellType");
masterTable = renamevars(masterTable,"Edit","FractionOfReadsEdited");

% Change CellID format to make single word string
cellidcol = string(masterTable.CellID);
newcellid = split(cellidcol,"-");
masterTable.CellID = append ("d",newcellid(:,2));

%% Save Imported data as matlab file
save([path.m 'masterTable.mat'],"masterTable");
```
