## Supplementary material for "A stochastic RNA editing process targets a select number of sites in individual *Drosophila* glutamatergic motoneurons": MATLAB code: Step2_Filter.m

```
% A stochastic RNA editing process targets a limited number of sites in individual Drosophila glutamatergic motoneurons
% Andrés B. Crane, Suresh K. Jetti, J. Troy Littleton
% The Picower Institute for Learning and Memory, Department of Brain and Cognitive Sciences, Department of Biology, Massachusetts Institute of Technology, Cambridge, MA 02139
% Correspondence and requests for materials should be addressed to J.T.L..


%% Set up workspace and environment
clear
clc
path.main = 'C:\Users\Littleton Lab\Documents\Andres C drive\RNA editing\GATK analysis - Muscle vs neurons\';
path.gatkoutput = [path.main '\GATK output\'];
path.graphs = [path.main '\Results\'];
temp = open([path.gatkoutput 'masterTable.mat']);
masterTable = temp.masterTable;
clear temp;

%% Create table variable to indicate for each edit which filters it passed
masterTable.Filter = zeros(size(masterTable,1),1);

%% 1. Filter by minimum DP
minDP = 10;
masterTable.Filter(masterTable.Depth>minDP) = 1;


%% 2. Filter by minimum qual
minQual = 20;
rows = (masterTable.Filter==1) & (masterTable.QUAL>minQual);
masterTable.Filter(rows) = 2;


%% 3. Filter by minimum edit percentage
minEdit = 0.1;
rows = (masterTable.Filter==2) & (masterTable.FractionOfReadsEdited>minEdit);
masterTable.Filter(rows) = 3;


%% 4. Filter by minimum cell number
minCellNum = 10;
rows = (masterTable.Filter==3) & (min_cells_with_reads_filter(masterTable,minCellNum));
masterTable.Filter(rows) = 4;


%% How many unique edit sites?
%num = unique(masterTable.('Edit Position'));

%% Save filtered edits
save([path.gatkoutput 'filtered edits'],"masterTable")

%% Graph A to G changes
graphFiltering(path,masterTable,minDP,minQual,minEdit,minCellNum);
%% Functions
function indeces_to_keep = min_cells_with_reads_filter(input,min_cells_with_reads)

    % Use only rows that passed previous filters
    rows = input.Filter>=3;

    % Save tables of just Ib and just Is rows
    input_Ib = input(rows & input.CellType == "Ib",:);
    input_Is = input(rows & input.CellType == "Is",:);
    input_M = input(rows & input.CellType == "M",:);

    % Count number of times each position appears
    Ib_groupcounts = groupcounts(input_Ib,"Edit Position");
    Is_groupcounts = groupcounts(input_Is,"Edit Position");
    M_groupcounts = groupcounts(input_M,"Edit Position");

    % Save the positions that occur more than "min_cell_with_reads" times
    Ib_Positions_over_min_cells = Ib_groupcounts(Ib_groupcounts.GroupCount > min_cells_with_reads,"Edit Position");
    Is_Positions_over_min_cells = Is_groupcounts(Is_groupcounts.GroupCount > min_cells_with_reads,"Edit Position");
    M_Positions_over_min_cells = M_groupcounts(M_groupcounts.GroupCount > min_cells_with_reads,"Edit Position");

    % Find out the indeces for the positions that pass the minimum
    % threshold. These make logicals
    Ib_indeces_to_keep = ismember(input.('Edit Position'),Ib_Positions_over_min_cells{:,1});
    Is_indeces_to_keep = ismember(input.('Edit Position'),Is_Positions_over_min_cells{:,1});
    M_indeces_to_keep = ismember(input.('Edit Position'),M_Positions_over_min_cells{:,1});

    % Use the indeces to save those rows out of the original input

    % !!!!! IMPORTANT !!!!!! If there is a position that passed the minimum
    % number of cells threshold in only Is, but there are some Ib cells
    % that also have editing at that same position (though not enough to
    % meet the minimum), BOTH the Ib and the Is indeces are kept. This
    % means that some Ib cells will be listed as edits, even though truly
    % it is the Is that has the editing. However, it is important to keep
    % the Ib quantification at this position for further processing later.

    %filtered_by_min_cell_with_reads = input(Ib_indeces_to_keep | Is_indeces_to_keep,:);
    indeces_to_keep = Ib_indeces_to_keep | Is_indeces_to_keep | M_indeces_to_keep;

end


function graphFiltering(path,masterTable,minDP,minQual,minEdit,minCellNum)

    f = figure(1);
    
    % Tiles
    tiledlayout(5,1);
    nexttile;
    
    
    changes = append(string(masterTable.REF),string(masterTable.ALT));
    histogram(categorical(changes))
    title('GATK output with SNPs, indels, and DP=0 removed')
    
    nexttile;
    rows = masterTable.Filter>=1;
    changes = append(string(masterTable.REF(rows)),string(masterTable.ALT(rows)));
    histogram(categorical(changes))
    title(['Filter 1: After min DP = ' num2str(minDP)])
    
    nexttile;
    rows = masterTable.Filter>=2;
    changes = append(string(masterTable.REF(rows)),string(masterTable.ALT(rows)));
    histogram(categorical(changes))
    title(['Filter 2: After min QUAL = ' num2str(minQual)])
    
    nexttile;
    rows = masterTable.Filter>=3;
    changes = append(string(masterTable.REF(rows)),string(masterTable.ALT(rows)));
    histogram(categorical(changes))
    title(['Filter 3: After min edit rate = ' num2str(minEdit)])
    
    nexttile;
    rows = masterTable.Filter>=4;
    changes = append(string(masterTable.REF(rows)),string(masterTable.ALT(rows)));
    histogram(categorical(changes))
    title(['Filter 4: After min cell number = ' num2str(minCellNum)])
    
    % Overall graph
    %title(f,'Filtering effect on base edit distribution')
    %xlabel(f,'Edit base change')
    %ylabel(f,'Number of occurences')
    
    exportgraphics(f,[path.graphs '\FilteringEffectOnBaseEditDistribution.pdf'])
end
```
