## Supplementary material for "A stochastic RNA editing process targets a select number of sites in individual *Drosophila* glutamatergic motoneurons": MATLAB code: Step3_Process.m

```
% A stochastic RNA editing process targets a limited number of sites in individual Drosophila glutamatergic motoneurons
% Andrés B. Crane, Suresh K. Jetti, J. Troy Littleton
% The Picower Institute for Learning and Memory, Department of Brain and Cognitive Sciences, Department of Biology, Massachusetts Institute of Technology, Cambridge, MA 02139
% Correspondence and requests for materials should be addressed to J.T.L..


%% Set up workspace and environment
clear
clc

path.main = 'C:\Users\Littleton Lab\Documents\Andres C drive\RNA editing\GATK analysis - Muscle vs neurons\';
path.gatkoutput = [path.main '\GATK output\'];
path.flybasedata = [path.main '\Flybase genomic data\'];
path.results = [path.main '\Results\Canonical'];
path.tables = [path.main '\Results\Canonical\Tables\'];
path.tpms = [path.main '\TPM data\'];

% Load filtered edits
temp = open([path.gatkoutput 'filtered edits.mat']);
data_edits = temp.masterTable;
clear temp;

% Load flybase data (create this using process_flybase_data.mat)
disp("Loading flybase data")
flybase_data = load([path.flybasedata 'flybase_data_2023-07-20_r6.51.mat']);   %contains start and end position for cds, 5 and 3' UTR, and sequence info from standard genome on flybase
flybase_data = flybase_data.flybase_data;
constantFlybase_data = parallel.pool.Constant(flybase_data);
constantFlybase_data = constantFlybase_data.Value;
disp("Loading flybase data...done")

 %% Only use edits from cells with ADAR expression detected
% runsection = true;
% 
% % Load TPMs
% if ~exist('tpmdata','var')
%     tpmdata = readtable([path.tpms 'Jetti et al - Supp Table 1.xlsx'],'Range','A5:JC15032');
% end
% 
% if runsection == true
% 
%     % Only use filtered edits
%     data_edits = data_edits(data_edits.Filter>=max(data_edits.Filter),:);
% 
%     % Add adar expression info to data_edits
%     data_edits = adar(data_edits,tpmdata);
% 
%     % Delete edits in cells with 0 adar expression
%     %data_edits = data_edits(data_edits.("ADAR TPM")>0,:);
% 
%     % Save edits in cells with 0 adar expression, delete the rest
%     data_edits = data_edits(data_edits.("ADAR TPM")==0,:);
% 
% else
%     data_edits = adar(data_edits,tpmdata);
% 
% end

%% Process filtered edits into a summary

data_summary = data_processing(data_edits);

%% Double check for genomic (DNA) SNPs using aligned reads instead of GATK


    % Creat list of genomic SNPs
    threshold = 0.96;   %if <= this fraction of DNA reads is the same, this site is labeled as a SNP
    genomicSNPs = findGenomicSNPs(data_summary, threshold);


    % Delete any genomic SNPs
    for i = 1:size(genomicSNPs,1)
        editPosition = genomicSNPs.("Edit Position")(i);

        idx_sum = data_summary.("Edit Position") == editPosition;
        idx_edi = data_edits.("Edit Position") == editPosition;

        data_summary(idx_sum,:) = [];
        data_edits(idx_edi,:) = [];
    end
    
    clear data i idx_sum idx_edi
%% From this point on only use data_edits that passed filters
data_edits = data_edits(data_edits.Filter>=max(data_edits.Filter),:);

%% quick fix
genomicSNPs97 = readtable('../Results/Genomic SNPs 3 percent threshold.xlsx');

genomicSNPs97.StillASNPat4Percent = repmat("-",size(genomicSNPs97,1),1);

for i = 1:size(genomicSNPs97,1)

    if sum(genomicSNPs97.('EditPosition')(i) == genomicSNPs.("Edit Position")) > 0
        genomicSNPs97.StillASNPat4Percent(i) = "Yes";
    end
end

writetable(genomicSNPs97,'../Results/Genomic SNPs comparison 3 and 4 percent.xlsx')

%% Add gene name and flybase ID
data_edits = addFlybaseID(data_edits,path);
data_summary = addFlybaseID(data_summary,path);

%% Add Gene part (CDS, UTR, etc) and edited codon
    data_summary = addGenePart(data_summary,constantFlybase_data);
    
    % Delete edits occuring in overlaping genes from non-exons in
    % data_edits
    OverlappingGenes = readtable('../Results/Overlapping Genes.xlsx');
    for i = 1:size(OverlappingGenes)
        editPosition = OverlappingGenes.EditPosition(i);
        idx_edit = data_edits.("Edit Position") == editPosition;
        data_edits(idx_edit,:) = [];
    end

clear i idx_edit
%% Add previously known RNA editing sites
data_summary = addKnownRNAeditSites(data_summary);

%% Add the transcripts per million RNA sequencing values for each gene and ADAR

% Load TPMs
if ~exist('tpmdata','var')
    tpmdata = readtable([path.tpms 'Jetti et al - Supp Table 1.xlsx'],'Range','A5:JC15032');
end

data_summary = TPMs(data_summary,tpmdata);
data_edits = TPMs(data_edits,tpmdata);
data_edits = adar(data_edits,tpmdata);

%% Add codon amino acid change from edit

data_summary = editedCodon(data_summary,constantFlybase_data);
data_summary = CDSmutationType(data_summary);

%% Remove edits that occur very close to the 5'UTR start site

indecesToRemove = data_summary.PosInGenePart < 20 ...
    & data_summary.GenePart == "FivePrimeUTR";
data_summary(indecesToRemove,:) = [];

%% Add Pfam domain
data_summary = addPfamDomain(data_summary);

%% Add best gene summary
data_summary = addGeneSummary(data_summary);

%% Check for neighboring edit sites and note distance from closest edit
data_summary = checkForNeighbors(data_summary);

%% Add read depth for each site
data_summary = coverage(data_summary, data_edits);

%% Mark edits as canonical (A-->G) or not and biological change
data_summary = markCanonical(data_summary);
data_edits = markCanonical(data_edits);

%% Add what cell types each edit occurs in
data_summary = addEditOccurrence(data_summary);

%% BLAST
%data_summary = myblast(data_summary);

%% Save data

% Rearrange variables in table for easier access
data_summary = movevars(data_summary,{'Edit Position',...
                                      'GenePart',...
                                      'BiologicalBaseChange',...
                                      'AminoAcid_Original',...
                                      'AminoAcid_Edited',...
                                      'Mean Ib editing',...
                                      'Mean Is editing',...
                                      'Mean M editing',...
                                      'MutationType'},'After','Gene');

% Canonical edits
writetable(data_summary(data_summary.Canonical==1,:),[path.tables 'Supplemental Table 2 — RNA Editing Summary for Canonical Edits.xlsx'],'WriteMode','overwrite');
writetable(data_edits(data_edits.Canonical==1,:),[path.tables 'Supplemental Table 1 — All Canonical Edits.xlsx'],'WriteMode','overwrite');

%Noncanonical edits
writetable(data_summary(data_summary.Canonical==0,:),[path.main '\Results\Noncanonical\Tables\' 'Supplemental Table 4 — RNA Editing Summary for Non-Canonical Edits.xlsx'],'WriteMode','overwrite');
writetable(data_edits(data_edits.Canonical==0,:),[path.main '\Results\Noncanonical\Tables\' 'Supplemental Table 3 — All Non-Canonical Edits.xlsx'],'WriteMode','overwrite');

save([path.main '\Results\' 'Matlab files'], 'data_summary','data_edits');
%% Functions
function genomicSNPs = findGenomicSNPs(data_summary, threshold)
    % Load the list of edit sites to check
    %load("../Results/Matlab files.mat")
    chromosome = data_summary.Chromosome;
    position = data_summary.Position;

    % Load DNA sequencing files
    addpath 'C:\Users\Littleton Lab\Documents\Andres C drive\RNA editing\2023-02-28 DNA sequencing of Ib, Is-gal4, and UAS-GFP by Duan Ma\BAM aligned with BWA'
    fprintf('Load DNA alignments........................................');
    load('DNA_Alignment_DuanMaBWA.mat')
    constantBioMap_DNA_Ib = parallel.pool.Constant(constantBioMap_DNA_Ib);
    constantBioMap_DNA_Is = parallel.pool.Constant(constantBioMap_DNA_Is);
    constantBioMap_DNA_GFP = parallel.pool.Constant(constantBioMap_DNA_GFP);
    fprintf('done\n');

    %% Initialize variables for parallel processing
    numGenes = size(chromosome,1);
    constantChromosome = parallel.pool.Constant(chromosome);
    constantPosition = parallel.pool.Constant(position);
    data = repmat('-',numGenes,3);
    percentDNA = nan(numGenes,3);  %percentDNA stores the percentage of reads with the most common nucleotide

    % Create a waitbar
    D = parallel.pool.DataQueue;
    afterEach(D, @nUpdateWaitbar);
    f = waitbar(0,'Finding potential edit sites');
    global progress;
    progress = 0;
    send(D,[1 numGenes]);

    parfor gene = 1:numGenes %go through each gene in the genelist 1:numGenes parfor loop
    
        % Get chrom, start, and end pos from gene list
        currentChromosome = constantChromosome.Value{gene};
        currentPosition = constantPosition.Value(gene);
        disp(currentPosition)
    
        % Get most common DNA nucleotide in the DNA sequence
        referenceDNA = repmat('-',1,3);   %referenceDNA = saves the maximum nucleotide in profile as a character
        percentDNArow = nan(1,3);
        alignment = char();
        BioMapList =  [constantBioMap_DNA_GFP.Value;...
                      constantBioMap_DNA_Ib.Value;...
                      constantBioMap_DNA_Is.Value];
            for i = 1:size(BioMapList,1)
                
                % Get compact alignment for gene from biomap
                alignment = getCompactAlignment(BioMapList(i), currentPosition, currentPosition, currentChromosome);
                % Record num of reads for each position
                %reads{i} = sum(isstrprop(alignment, "alpha"));
        
                % Change spaces into dashes
                alignment = strrep(string(alignment),' ','-');
        
                % Profile outputs 4 by x array of percentage occurences in this order: 4 nucleotides — A C G T/U
                %profiles{i} = seqprofile(alignment,'Alphabet','n');
                profile = seqprofile(alignment,'Alphabet','n');
    
                % Find the maximum value nucleotide for each position in the
                % profile
    
                [M1,I1] = max(profile);
                
                if size(M1,1)>0 && size(M1,2)>0
                    percentDNArow(1,i) = M1;
                end
    
                % We want to find genomic SNPs where it might not be present
                % fully within a single fly population, but allow enough wiggle
                % room to not catch sequencing errors
    
                if M1 > threshold
                    % ex if 97% of the DNA is a single nucleotide, the remaining
                    % 3% are probably sequencing error or can be safely ignored
    
                    % Save character representation of max nuc
                    referenceDNA(I1==1,i) = 'A';
                    referenceDNA(I1==2,i) = 'C';
                    referenceDNA(I1==3,i) = 'G';
                    referenceDNA(I1==4,i) = 'T';
                else
                    % If <= 97% of the DNA is the same, we will flag it
                    % as a potential genomic SNP
    
                    referenceDNA(1,i) = '-';
                end
    
    
    
            end
    
        percentDNA(gene,:) = percentDNArow;
        data(gene,:) = referenceDNA;
        send(D,[1 numGenes]);
    end

    
    %% Double check if any are genomic SNPs where all 3 DNAs do not match
    
    % Make an index of SNPs where our fly DNA does not match flybase ref DNA
    indexSNPs = (char(data_summary.REF) ~= data(:,1) | ...
                       char(data_summary.REF) ~= data(:,2) | ...
                       char(data_summary.REF) ~= data(:,3));
    genomicSNPs = data_summary(indexSNPs,:);
    
    %% SAVE matlab format data
    save('../Results/doubleCheckForGenomicSNPs','data', 'genomicSNPs',"percentDNA")

    %% Save new spreadsheet file with genomic snps found
    writetable(genomicSNPs,'../Results/Genomic SNPs.xlsx','WriteMode','overwrite')
end

function input = myblast(input)

    % Number of sequences to BLAST
    numseq = size(input,1);

    % For each sequence, generate a request ID from the BLAST server
    RID = strings(numseq,1);
    RTOE = [];
    orgs = ['Mus musculus[organism] OR '...
            'Homo sapiens[organism] OR '...
            'Aedes aegypti[organism]'];
    for i = 1:numseq
        seq = input.AminoAcid_Sequence_align{i};

        if isempty(seq)==1
            continue
        end

        [RID(i), RTOE(i)] = blastncbi(seq,'blastp','Entrez',orgs);
    end
    
    % Wait until the longest job is done
    pause(max(RTOE)*60);

    % Make table to hold results
    PerID = table();    %percent identity to query sequence


    % For each RID, download the results from the NCBI server
    blastdata = getblast(RID(4));
    numHits = size(blastdata(:).Hits,2);

        % Extract the species name
        sp=strings(numHits,1);
        for i = 1:numHits
            def = blastdata.Hits(i).Definition;
            ext = extractBetween(def,'[',']');
            sp(i,1) = ext(1);
        end


        % Calculate percent identity
        id = [];
        for i = 1:numHits
            id(i,1) = blastdata.Hits(i).Hsps(1).Identities ./ ...
                blastdata.Hits(i).Hsps(1).AlignmentLength * 100;
        end       


        % For each species name, calculate the max percent identity
        index.hs = sp == "Homo sapiens";
        index.mm = sp == "Mus musculus";
        index.ce = sp == "Caenorhabditis elegans";
        index.aa = sp == "Aedes aegypti";


        PerID.Human = max(id(index.hs));
        PerID.Mouse = max(id(index.mm));
        perID.Celegans = max(id(index.ce));
        perID.Mosquito = max(id(index.aa));


end

function input = addEditOccurrence(input)

% Find which edits are found in more than 10 samples of Ib, Is, or M
    IndexNeuronEdits = (input.("Num of Ib cells with edit") > 10)...
                     | (input.("Num of Is cells with Edit") > 10);
    
    IndexMuscleEdits = (input.("Num of M samples with edit") > 10);

% Record N or M (or both) in new column
    neuronEdits = strings(size(input,1),1);
    muscleEdits = strings(size(input,1),1);

    neuronEdits(IndexNeuronEdits) = "N";
    muscleEdits(IndexMuscleEdits) = "M";

    input.("Cell types with edit") = strcat(neuronEdits,muscleEdits);
end

function input = adar(input,tpmdata)

    % Find ADAR in tpm data and extract
    index = strcmp('FBgn0026086',tpmdata.FlybaseID);
    adarTPMs = [tpmdata(index,12:116) ...   %Ib cells
                tpmdata(index,118:218)...   %Is cells
                tpmdata(index,220:236)...   %M1 cells
                tpmdata(index,238:263)];    %M4 cells
    
    tableColnames = adarTPMs.Properties.VariableNames;
    numOfEdits = size(input,1);
    adarTPMsAccum = nan(numOfEdits,1);
    % For each RNA edit entry row in input
    for i = 1:numOfEdits

        % Find the cell name in adar tpm table
        index = contains(tableColnames,input.CellID(i),'IgnoreCase',true);

        % Extract TPM for that cell
        adarTPMsAccum(i) = adarTPMs{1,index};

        if mod(i,1000)==0
            disp(append("Adding ADAR: ",num2str(i)))
        end
    end

    % Add TPM info to input table
    input.("ADAR TPM") = adarTPMsAccum;

%%TEMP
% temp=table;
% for i = 1:size(adarTPMs,2)
% 
%     if adarTPMs{1,i} ~= 0
%         temp.cell(i) = tableColnames(1,i);
%         temp.adar(i) = adarTPMs{1,i};
%     end
% 
% end
% 
% for i = 1:size(temp,1)
%     idx(i) = isempty(temp.cell{i});
% end
% temp(idx,:) = [];
% 
% mean(temp.adar(61:end))


end

function input = TPMs(input,tpmdata)

    numOfEdits = size(input,1);
    tpm_Ib = nan(numOfEdits,1);
    tpm_Is = nan(numOfEdits,1);
    pval = nan(numOfEdits,1);
    tpm_M1 = nan(numOfEdits,1);
    tpm_M4 = nan(numOfEdits,1);
    % For each RNA edit entry row in input
    for i = 1:numOfEdits

        % find the row number of that gene in tpmdata
        index = strcmp(input.FlybaseID(i),tpmdata.FlybaseID);

        % Check to make sure an index is found, if not, skip
        if sum(index)~=1
            continue
        end

        % Extract the Ib and Is average TPM expression and significance
        tpm_Ib(i) = tpmdata.IbTPMsAverage(index);
        tpm_Is(i) = tpmdata.IsTPMsAverage(index);
        pval(i) = tpmdata.Ib_IsAdjustedP_Value(index);
        tpm_M1(i) = tpmdata.M1TPMsAverage(index);
        tpm_M4(i) = tpmdata.M4TPMsAverage(index);
        
        if mod(i,1000)==0
            disp(append("Adding TPMS: ",num2str(i)))
        end
    end

    % Add TPM info to input table
    input.("Gene TPM in Ib") = tpm_Ib;
    input.("Gene TPM in Is") = tpm_Is;
    input.("Gene TPM adj pval") = pval;
    input.("Gene TPM in M1") = tpm_M1;
    input.("Gene TPM in M4") = tpm_M4;

    if ismember('GenePart', input.Properties.VariableNames)
        input = movevars(input,("Gene TPM in Ib"),'After','GenePart');
        input = movevars(input,("Gene TPM in Is"),'After',("Gene TPM in Ib"));
        input = movevars(input,("Gene TPM adj pval"),'After',("Gene TPM in Is"));
        input = movevars(input,("Gene TPM in M1"),'After',("Gene TPM adj pval"));
        input = movevars(input,("Gene TPM in M4"),'After',("Gene TPM in M1"));
    end
end

function input = CDSmutationType(input)

    % Create variables to be added to input table
    MutationType = repmat("-",size(input,1),1);
    MutationType_AA_group_change = repmat("-",size(input,1),1);

    % Extract all the CDS rows
    idx_cds = strcmp(input.GenePart,'CDS');
    original = string(input.AminoAcid_Original(idx_cds));
    edited = string(input.AminoAcid_Edited(idx_cds));

    % Create variables to hold AA change just for CDS rows
    MutationTypeCDS = repmat("-",size(original,1),1);
    MutationTypeCDS_AA_group_change   = repmat("-",size(original,1),1);
    


    % Find types of mutations
    idx_nonsense = strcmp('*',edited);
    MutationTypeCDS(idx_nonsense,1) = "Nonsense";
    
    idx_silent = strcmp(original,edited);
    MutationTypeCDS(idx_silent,1) = "Silent";

    idx_missense = and(~idx_silent,~idx_nonsense);
    MutationTypeCDS(idx_missense,1) = "Missense";
    
    % Save MutationTypeCDS into MutationType and into table variable
    MutationType(idx_cds,1) = MutationTypeCDS;
    input.MutationType = MutationType;


    % Find when the AA changes groups (polar to non-polar, etc)
    original_missense = original(idx_missense);
    edited_missense = edited(idx_missense);
    groupchange = repmat("No",size(original_missense));

    AA_group_original = determineAAGroup(original_missense);
    AA_group_edited = determineAAGroup(edited_missense);

    % Save into table variable
    idx_groupchange = AA_group_original~=AA_group_edited;
    groupchange(idx_groupchange) = "Yes";
    MutationTypeCDS_AA_group_change(idx_missense) = groupchange;
    MutationType_AA_group_change(idx_cds,1) = MutationTypeCDS_AA_group_change;
    input.AminoAcidGroupChange = MutationType_AA_group_change;

    % Arrange table variables order
    input = movevars(input,["MutationType" "AminoAcid_Original" ...
        "AminoAcid_Edited" "AminoAcid_Sequence"],'After','GenePart');

    function output = determineAAGroup(input)
        output = zeros(size(input));
        group={};
        group{1,1} = {'R','K'};
        group{2,1} = {'D','E'};
        group{3,1} = {'S','T'};
        group{4,1} = {'N','Q'};
        group{5,1} = {'A','V','I','L','M','F','Y','W'};
        group{6,1} = {'C'};
        group{7,1} = {'G'};
        group{8,1} = {'P'};
        group{9,1} = {'H'};

        for i = 1:9
            idx = ismember(input,group{i});
            output(idx) = i;
        end

    end
end

function input = addKnownRNAeditSites(input)
   
    
    % Manually download RNA editing glyphs from jbrowse by enabling Transcript
    % Level Features > RNA editing sites, then clicking the down arrow on the
    % track name >Save track data > Whole reference sequence as gff3 file. Do
    % this for each chromosome
    
    % Load the gff3 files as GFFannotation objects
    path = "C:\Users\Littleton Lab\Documents\Andres C drive\RNA editing\Flybase genomic data\Previously known RNA editing sites\2023-04-17 RNA editing sites from flybase jbrowse";
    files = dir([char(path) '\*.gff3']);
    
    % Declare variables
    RNAeditingToAdd = string();               %temp variable inside of loop
    RNA_editing_sites_jbrowse = string();     %storage of all edit events

    % Open each gff file and accumulate master list of RNA editing events
    for i = 1:size(files,1)

        % Open the GFF file
        currentGFF = GFFAnnotation([files(i).folder '\' files(i).name]);
        numOfEdits = size(currentGFF.Start,1);

        % Save the chromosome name and nucleotide position of each RNA
        % editing event
        RNAeditingToAdd(1:numOfEdits,1) = string(currentGFF.Reference);
        RNAeditingToAdd(1:numOfEdits,2) = string(currentGFF.Start);

        % Accumulate a master list of all RNA editing events
        if RNA_editing_sites_jbrowse==""
            RNA_editing_sites_jbrowse = RNAeditingToAdd;
        else
            RNA_editing_sites_jbrowse = [RNA_editing_sites_jbrowse; RNAeditingToAdd];
        end

        % Clear variables
        clear RNAeditingToAdd
    end

    % Extract chr and position info from data_summary
    dsinfo = [string(input.Chromosome) string(input.Position)];
    

    % Find indeces in data_summary which match chr and position to RNA_editing_sites_jbrowse
    indeces = ismember(dsinfo,RNA_editing_sites_jbrowse,'rows');

    % Create new table variable for known editing sites
    input.PreviouslyKnownRNAeditingSite = repmat("No",size(input,1),1);
    input{indeces,"PreviouslyKnownRNAeditingSite"} = "Yes";
    input = movevars(input,"PreviouslyKnownRNAeditingSite",'After',"Edit Position");
end

function input = addPfamDomain(input)
   
    
    % Manually download pfam glyphs from jbrowse by enabling protein
    % level features > Protein Domains, then clicking the down arrow on the
    % Pfamtrack name >Save track data > Whole reference sequence as gff3 file. Do
    % this for each chromosome
    
    % Load the gff3 files as GFFannotation objects
    path = "C:\Users\Littleton Lab\Documents\Andres C drive\RNA editing\Flybase genomic data\Pfam protein domains";
    files = dir([char(path) '\*.gff3']);
    
    % Declare variables
    PfamDomains = table();      %storage of all edit events

    % Open each gff file and accumulate combined chr list of Pfam domains
    for i = 1:size(files,1)

        PfamDomainsTemp = table();   %temp variable inside of loop

        % Open the GFF file
        currentGFF = GFFAnnotation([files(i).folder '\' files(i).name]);

        % Save the chromosome name and nucleotide position of each Pfam
        % domain
        PfamDomainsTemp.Start = currentGFF.Start;
        PfamDomainsTemp.Stop = currentGFF.Stop;
        PfamDomainsTemp.Chr = currentGFF.Reference;
        PfamDomainsTemp.Strand = currentGFF.Strand;

        % Split the Attribute field to get the domain abbreviation info
        attr = currentGFF.Attributes;
        PfamDomainsTemp.PfamDomain = extractBetween(attr,"_Domain_name_abbreviation=",";");


        % Save into accumulation variable
        if isempty(PfamDomains)
            PfamDomains = PfamDomainsTemp;
        else
            PfamDomains = [PfamDomains; PfamDomainsTemp];
        end

        % Clear variables
        clear PfamDomainsTemp attr currentGFF
    end

    % For each rna edit in data_summary, find whether it matches to a
    % pfam domain between start and end positions, and record matches
    
    input.PfamDomain = repmat("-",size(input,1),1);
    input = movevars(input,"PfamDomain","After","MutationType");

    for i = 1:size(input,1)

        % If the current position is not CDS, then don't look for pfam
        if input.GenePart(i) ~= "CDS"
            continue
        end
    
        % Find indeces in PfamDomain for current chr and position
        index.chr = strcmp(PfamDomains.Chr,input{i,"Chromosome"});
        index.pos = (PfamDomains.Start <= input{i,"Position"}) ...
                  & (input{i,"Position"} <= PfamDomains.Stop);
        index.match = index.chr & index.pos;

        % If there is no domain here, continue
        domain = PfamDomains{find(index.match,1),"PfamDomain"};
        if isempty(domain)
            continue
        end

        % Record domain into variable
        input.PfamDomain(i) = domain;

        % Tracking
        if mod(i,100)==0
        disp(i)
        end
    end

end

function input = addGenePart(input,constantFlybase_data)
    %% Add reference DNA info
    
    % Load flybase data
    f = waitbar(0,'Starting...');
    
    % Create a waitbar
    D = parallel.pool.DataQueue;
    afterEach(D, @nUpdateWaitbar);
    waitbar(0,f,'Add gene parts');
    global progress;
    progress = 0;
    send(D,[progress size(input,1)]);

    % Split location temporarily
    pos = string(input.("Edit Position"));
    temp = split(pos,":");
    rna.chr = categorical(temp(:,1));
    rna.pos = str2double(temp(:,2));
    clear temp pos
    
    
    % Initialize variable GenePart to hold "cds" "utr" etc
    GenePart = repmat("-",[size(input,1) 1]);
    PosInGenePart = nan(size(input,1),1);
    PosInGenePartFrac = nan(size(input,1),1);
    
    % Go through each position (parfor loop)
    for position = 1:size(input,1)
    
        
        % Find the gene in flybase_data
        indeces_fbd = strcmp([constantFlybase_data.GeneID],input.FlybaseID(position));
    
        if sum(indeces_fbd)==0
             GenePart(position) = "NotAnnotatedInFlyBase";
            continue
        end
    
        % Turn indeces into column subscripts of where the transcripts are
        % in flybase_data
        flybasetranscripts = find(indeces_fbd);
        genepartlist = [];
        posInGenePartAccum = [];
        posInGenePartFracAccum = [];
        editPosition = rna.pos(position);
    
        % For each transcript
        for transcript = 1:size(flybasetranscripts,2)
            
            % Get gene parts and see if editPosition is in them
            try
                cds = constantFlybase_data(flybasetranscripts(transcript)).CDS;
                cds_mat = cell2mat(cds(:,1:2));
                cds_yn = (cds_mat(:,1) <= editPosition) &  ( editPosition <= cds_mat(:,2));
            catch
            end
    
            try
                utr5 = constantFlybase_data(flybasetranscripts(transcript)).FivePrimeUTR(:,1:2);
                utr5 = cell2mat(utr5);
                utr5_yn = (utr5(:,1) <= editPosition) &  ( editPosition <= utr5(:,2));
                utr5_nucsFromStart = nan;
                utr5_nucsFromStartFrac = nan;
                
                % If current edit is in FivePrimeUTR, calculate distance
                % from start
                if sum(utr5_yn)>0
                    utr5_totalLength = sum(utr5(:,2)-utr5(:,1));
                    pos = find(utr5_yn);
                    
        
                    % plus strand genes
                    if constantFlybase_data(flybasetranscripts(transcript)).DNAstrand == "+"
                        utr5(utr5_yn,2) = editPosition;
                        utr5_nucsFromStart = utr5(1:pos,2)-utr5(1:pos,1);
                    
                    % minus strand genes
                    elseif constantFlybase_data(flybasetranscripts(transcript)).DNAstrand == "-"
                        utr5(utr5_yn,1) = editPosition;
                        pos = size(utr5_yn,1)-pos;
                        utr5_nucsFromStart = utr5(end-pos:end,2)-utr5(end-pos:end,1);
                    end
        
                    utr5_nucsFromStartFrac = utr5_nucsFromStart./utr5_totalLength;
                end
            catch
            end
    
            try
                utr3 = constantFlybase_data(flybasetranscripts(transcript)).ThreePrimeUTR(:,1:2);
                utr3 = cell2mat(utr3);
                utr3_yn = (utr3(:,1) <= editPosition) &  ( editPosition <= utr3(:,2)); 
    
                utr3_totalLength = sum(utr3(:,2)-utr3(:,1));
                pos = find(utr3_yn);
                utr3(utr3_yn,2) = editPosition;
    
                % plus strand genes
                if constantFlybase_data(flybasetranscripts(transcript)).DNAstrand == "+"
                    utr3_nucsFromStart = utr3(1:pos,2)-utr3(1:pos,1);
                
                % minus strand genes
                elseif constantFlybase_data(flybasetranscripts(transcript)).DNAstrand == "-"
                    pos = size(utr3_yn,1)-pos;
                    utr3_nucsFromStart = utr3(end-pos:end,2)-utr3(end-pos:end,1);
                end
    
                utr3_nucsFromStartFrac = utr3_nucsFromStart./utr3_totalLength;
            catch
            end
    
            % Accumulate the gene part in a list
            if sum(cds_yn)>0
                genepartlist = [genepartlist;1];
            elseif  sum(utr5_yn)>0
                genepartlist = [genepartlist;2];
                posInGenePartAccum = [posInGenePartAccum;utr5_nucsFromStart];
                posInGenePartFracAccum = [posInGenePartFracAccum;utr5_nucsFromStartFrac];
            elseif  sum(utr3_yn)>0
                genepartlist = [genepartlist;3];
                posInGenePartAccum = [posInGenePartAccum;utr3_nucsFromStart];
                posInGenePartFracAccum = [posInGenePartFracAccum;utr3_nucsFromStartFrac];
            end
        end
    
        % Find median genepartlist and save it
         mediangenepart = median(genepartlist);
    
         if mediangenepart == 1
            GenePart(position) = "CDS";
         elseif mediangenepart == 2
            GenePart(position) = "FivePrimeUTR";
            PosInGenePart(position) = median(posInGenePartAccum);
            PosInGenePartFrac(position) = median(posInGenePartFracAccum);
         elseif mediangenepart == 3
            GenePart(position) = "ThreePrimeUTR";
            PosInGenePart(position) = median(posInGenePartAccum);
            PosInGenePartFrac(position) = median(posInGenePartFracAccum);
         end
    
         % Update waitbar
        send(D, [position,size(input,1)]);
    
    
    
    end
    
    % Add GeneParts to data_summary
    input = addvars(input,GenePart,'After','Edit Position');
    input.PosInGenePart = PosInGenePart;
    input.PosInGenePartFrac = PosInGenePartFrac;

    % Identify rows that did not match to a genePart (these are where genes
    % overlap, but is in an intron in this gene but an actual genePart in
    % another gene)
    indeces = strcmp(input.GenePart,"-") | strcmp(input.GenePart,"NotAnnotatedInFlyBase");

    % Save table of which rows were deleted. Use this to delete the same
    % entries in data_edits
    writetable(input(indeces,:),'../Results/Overlapping Genes.xlsx','WriteMode','overwrite')

    % Delete edits occuring in overlapping gene parts
    input(indeces,:) = [];
    close(f);
end

function input = editedCodon(input,constantFlybase_data)


    
    % Load flybase data
    f = waitbar(0,'Loading flybase data');
    constantFlybase_data = rmfield(constantFlybase_data,{'FivePrimeUTR','ThreePrimeUTR'});

    
    % Find positions that are labeled CDS
    CDSpositionsInInput = find(strcmp(input.GenePart,'CDS'));

    % Create a waitbar
    D = parallel.pool.DataQueue;
    afterEach(D, @nUpdateWaitbar);
    waitbar(0,f,'Adding codon info');
    global progress;
    progress = 0;
    send(D,[progress size(CDSpositionsInInput,1)]);
    

    % Initialize variables to hold codon and amino acid
    input.Codon_Original = repmat({'---'},size(input,1),1);
    input.Codon_Edited = repmat({'---'},size(input,1),1);
    input.AminoAcid_Original = repmat('-',size(input,1),1);
    input.AminoAcid_Edited = repmat('-',size(input,1),1);


    % Go through each position (parfor loop)
    parfor position = 1:size(CDSpositionsInInput,1)

        currentPos = CDSpositionsInInput(position);
        found = false;

        % Find the gene in flybase_data
        indeces_fbd = strcmp([constantFlybase_data.GeneID],input.FlybaseID(currentPos));
    
        if isempty(indeces_fbd)
            continue
        end

        % Turn indeces into column subscripts of where the transcripts are
        % in flybase_data
        flybasetranscripts = find(indeces_fbd);
        editPosition = input.Position(currentPos);

    
        % For each transcript
        for transcript = 1:size(flybasetranscripts,2)

            if found == true
                break
            end

            % Determine if edit is contained in this transcript's CDS
            cds = constantFlybase_data(flybasetranscripts(transcript)).CDS;
            cds_mat = cell2mat(cds(:,1:2));
            cds_yn = (cds_mat(:,1) <= editPosition) &  ( editPosition <= cds_mat(:,2));

            % If this particular transcript does not have a CDS at this
            % location, continue to the next
            if sum(cds_yn)==0
                continue
            end
            
            % Find if the gene is on + or - DNA strand (affects codon)
            currentDNAStrand = constantFlybase_data(flybasetranscripts(transcript)).DNAstrand;  


            % Find the genomic position number of the first nucleotide in
            % the codon
            [editRegionNucsShort,editRegionNucsLong,modNumNucleotides,~] = findFirstNucOfCodon(currentDNAStrand,cds_mat,cds_yn,editPosition,cds);
     
            % Calculate edited codon
            try
            codon = editRegionNucsShort(1:3);
            catch
                disp(["exit at transcript " num2str(transcript) " position " num2str(currentPos)])
            end
            codon_edited = codon;

            if currentDNAStrand == "+"
                switch modNumNucleotides
                    case 0
                        codon_edited(3) = char(input.ALT(currentPos));
                    case 2
                        codon_edited(2) = char(input.ALT(currentPos));
                    case 1
                        codon_edited(1) = char(input.ALT(currentPos));
                end
            else
                switch modNumNucleotides
                    case 0
                        codon_edited(3) = seqcomplement(char(input.ALT(currentPos)));
                    case 2
                        codon_edited(2) = seqcomplement(char(input.ALT(currentPos)));
                    case 1
                        codon_edited(1) = seqcomplement(char(input.ALT(currentPos)));
                end
            end


            % Save codon and aa
            Codon_Original{position} = codon;
            Codon_Edited(position) = {codon_edited};
            AminoAcid_Original(position) = nt2aa(codon);
            AminoAcid_Edited(position) = nt2aa(codon_edited);
            
            found = true;
            AminoAcid_sequence{position} = nt2aa(editRegionNucsShort);
            AminoAcid_sequence_align{position} = nt2aa(editRegionNucsLong);
        end
    

    
         % Update waitbar
        send(D, [position,size(CDSpositionsInInput,1)]);
    
    %catch
    %warning(join(["Position = " string(position)]))
    %end
    end
    
    % Add Codon and Amino Acid to data_summary
    input{CDSpositionsInInput,'Codon_Original'} = Codon_Original';
    input{CDSpositionsInInput,'Codon_Edited'} = Codon_Edited';
    input{CDSpositionsInInput,'AminoAcid_Original'} = AminoAcid_Original';
    input{CDSpositionsInInput,'AminoAcid_Edited'} = AminoAcid_Edited';
    input{CDSpositionsInInput,'AminoAcid_Sequence'} = AminoAcid_sequence';
    input{CDSpositionsInInput,'AminoAcid_Sequence_align'} = AminoAcid_sequence_align';
    close(f);
end

function [editRegionNucsShort,editRegionNucsLong,modNumNucleotides,numNucleotides] = findFirstNucOfCodon(currentDNAStrand,cds_mat,cds_yn,editPosition,cds)
   % Determine num of nucleotides in seg containing position
    cds_mat(:,3) = zeros(size(cds_mat,1),1);


    % Make index for every seg up to seg containing position
    switch currentDNAStrand
        case '+'
            % For + strand genes, count every DNA segment from 1 to edit
            cds_mat(cds_yn,3) = editPosition - cds_mat(cds_yn,1)+1;
            cds_upto = logical(1:find(cds_yn)-1);
        case '-'
            % For - strand genes, count from last segment to edit
            cds_mat(cds_yn,3) = cds_mat(cds_yn,2) + 1 - editPosition;
            cds_upto = ones(size(cds_yn));
            cds_upto(1:find(cds_yn)) = zeros(size(1:find(cds_yn)));
            cds_upto = logical(cds_upto);
    end

    % For every seg up to seg containing position, subtract nucleotides
    if ~isempty(cds_upto)
        cds_mat(cds_upto,3) = cds_mat(cds_upto,2)-cds_mat(cds_upto,1)+1;
    end

    % Sum up num of nucleotides from beginning of CDS to edit
    numNucleotides = sum(cds_mat(:,3));

    % Find phase
    modNumNucleotides = mod(numNucleotides,3);


    % Combine bases into full length mRNA CDS sequence
    switch currentDNAStrand
        case '+'
            fullCDSseq = [cds{:,3}];
        case '-'
            fullCDSseq = [cds{:,3}];
    end
    

    % Extract codon + 27 more bases afterward if possible

    extraBases = 27;    %num of nucleotides to extract after edit codon
    basesTilEndOfCDS = size(fullCDSseq,2)-numNucleotides;

    if basesTilEndOfCDS < extraBases
        extraBases = floor(basesTilEndOfCDS/3)*3;   %make sure multiple of 3
    end

    switch modNumNucleotides
        case 1
            editRegionNucsShort = fullCDSseq(numNucleotides:numNucleotides+2+extraBases);
        case 2
            editRegionNucsShort = fullCDSseq(numNucleotides-1:numNucleotides-1+2+extraBases);
        case 0
            editRegionNucsShort = fullCDSseq(numNucleotides-2:numNucleotides-2+2+extraBases);
    end

    % Extract codon -90 and + 90 more bases  if possible

    extraBasesAfter = 90;    %num of nucleotides to extract after edit codon
    extraBasesBefore = 90;
    basesTilEndOfCDS = size(fullCDSseq,2)-numNucleotides;

    if basesTilEndOfCDS < extraBasesAfter
        extraBasesAfter = floor(basesTilEndOfCDS/3)*3;   %make sure multiple of 3
    end

    if numNucleotides <= extraBasesBefore
        extraBasesBefore = floor(numNucleotides/3)*3 - 3;   %make sure multiple of 3
    end

    switch modNumNucleotides
        case 1
            editRegionNucsLong = fullCDSseq(numNucleotides-extraBasesBefore:numNucleotides+2+extraBasesAfter);
        case 2
            editRegionNucsLong = fullCDSseq(numNucleotides-1-extraBasesBefore:numNucleotides-1+2+extraBasesAfter);
        case 0
            editRegionNucsLong = fullCDSseq(numNucleotides-2-extraBasesBefore:numNucleotides-2+2+extraBasesAfter);
    end


end

function nUpdateWaitbar(input)
    global progress;
    progress = progress + 1;
    waitbar(progress/input(2))
end

function data_processed = data_processing(input)

    disp("Starting data processing")

    % Split data into rows that passed filters and those that did not
    maxFilter = max(input.Filter);
    input_filterpass = input(input.Filter>=maxFilter,:);
    input_filterfail = input(input.Filter<maxFilter,:);

    % Find Ib and Is indeces
    IbcellIndex = input_filterpass.CellType == "Ib";
    IscellIndex = input_filterpass.CellType == "Is";
    McellIndex = input_filterpass.CellType == "M";

    % Find unique positions
    [g,ia,~] = unique(input_filterpass.('Edit Position'));

    % Build the stats table to summarize number and percent of edits
    stats = zeros(size(g,1),9);
    
    percentEdited_filterpass = input_filterpass.FractionOfReadsEdited;
    
    disp("Begin counting edits that passed filters")
    % Column 1: Edit position
    %------
    % Column 2: # of Ib cells with an edit at this position
    % Column 3: mean reads in Ib cells
    % Column 4: mean of all Ib cells' edit percent
    % Column 5: std dev of all Ib cells' edit percent
    %------
    % Column 6: # of Is cells with an edit at this position
    % Column 7: mean reads in Is cells
    % Column 8: mean of all Is cells' edit percent
    % Column 9: std dev of all Is cells' edit percent

    % Go through each unique location to get info about num and % of edits
    for i = 1:size(g,1)
        
        % Make an index for Ibs and Is at this position
        Ib = (input_filterpass.('Edit Position')==g(i)) & IbcellIndex;
        Is = (input_filterpass.('Edit Position')==g(i)) & IscellIndex;
        M = (input_filterpass.('Edit Position')==g(i)) & McellIndex;

        % Get stats
        stats(i,2)  = sum(Ib);
        stats(i,3)  = mean(input_filterpass.Depth(Ib));
        stats(i,4)  = mean(percentEdited_filterpass(Ib));
        stats(i,5)  = std(percentEdited_filterpass(Ib))/sqrt(length(percentEdited_filterpass(Ib)));
        stats(i,6)  = sum(Is);
        stats(i,7)  = mean(input_filterpass.Depth(Is));
        stats(i,8)  = mean(percentEdited_filterpass(Is));
        stats(i,9)  = std(percentEdited_filterpass(Is))/sqrt(length(percentEdited_filterpass(Is)));
        [~,p] = ttest2(percentEdited_filterpass(Ib),percentEdited_filterpass(Is));
        stats(i,10) = p;
        stats(i,11)  = sum(M);
        stats(i,12)  = mean(input_filterpass.Depth(M));
        stats(i,13)  = mean(percentEdited_filterpass(M));
        stats(i,14)  = std(percentEdited_filterpass(M))/sqrt(length(percentEdited_filterpass(M)));
        [~,p] = ttest2(percentEdited_filterpass(Ib),percentEdited_filterpass(M));
        stats(i,15) = p;
        [~,p] = ttest2(percentEdited_filterpass(Is),percentEdited_filterpass(M));
        stats(i,16) = p;

    end

    % Find NaN and convert to 0s
    stats(isnan(stats)) = 0;

    % Calculate difference between edit %
    stats(:,17) = stats(:,4) - stats(:,8);  %Ib - Is
    stats(:,18) = stats(:,4) - stats(:,12); %Ib - M
    stats(:,19) = stats(:,8) - stats(:,12); %Is - M

    % Calculate difference between num of cells
    stats(:,20) = stats(:,2) - stats(:,6);  %Ib - Is
    stats(:,21) = stats(:,2) + stats(:,6);  %Ib + Is
    stats(:,22) = stats(:,2) - stats(:,11); %Ib - M

    % Convert to table
    varNames = ["Edit Position",...
            "Num of Ib cells with edit",...
            "Mean reads in Ib cells with edit",...
            "Mean Ib editing",...
            "Std Dev of Ib editing",...
            "Num of Is cells with Edit",...
            "Mean reads in Is cells with edit",...
            "Mean Is editing",...
            "Std Dev of Is editing",...
            "P-val of Ib vs Is editing",...
            "Num of M samples with edit",...
            "Mean reads in M samples with edit",...
            "Mean M editing",...
            "Std Dev of M editing",...
            "P-val of Ib vs M editing",...
            "P-val of Is vs M editing",...
            "Mean Ib editing - mean Is editing",...
            "Mean Ib editing - mean M editing",...
            "Mean Is editing - mean M editing",...
            "Num of Ib cells - num of Is cells",...
            "Num of Ib cells + num of Is cells",...
            "Num of Ib cells - num of M cells"];
    data_processed = array2table(stats,'VariableNames',varNames);
    
    % Add the Location
    data_processed.("Edit Position") = g;
    data_processed.Chromosome = input_filterpass.CHROM(ia);
    data_processed.Position = input_filterpass.POS(ia);

    % Add the reference DNA and alt RNA
    REF = input_filterpass.REF(ia);
    ALT = input_filterpass.ALT(ia);

    data_processed = addvars(data_processed,REF,'After','Edit Position');
    data_processed = addvars(data_processed,ALT,'After','REF');

    % --------

    disp("Begin counting edits that did not pass filters")
    % For each position, calculate rate of editing in cells that did not
    % pass filters
    percentEdited_filterfail = input_filterfail.FractionOfReadsEdited;
    IbcellIndex = input_filterfail.CellType == "Ib";
    IscellIndex = input_filterfail.CellType == "Is";
    stats = zeros(size(g,1),8); 
    for i = 1:size(g,1)
        
        % Make an index for Ibs and Is at this position
        Ib = (input_filterfail.('Edit Position')==g(i)) & IbcellIndex;
        Is = (input_filterfail.('Edit Position')==g(i)) & IscellIndex;

        % Get stats
        stats(i,1)  = sum(Ib);
        stats(i,2)  = mean(input_filterfail.Depth(Ib));
        stats(i,3)  = mean(percentEdited_filterfail(Ib));
        stats(i,4)  = std(percentEdited_filterfail(Ib))/sqrt(length(percentEdited_filterfail(Ib)));
        stats(i,5)  = sum(Is);
        stats(i,6)  = mean(input_filterfail.Depth(Is));
        stats(i,7)  = mean(percentEdited_filterfail(Is));
        stats(i,8)  = std(percentEdited_filterfail(Is))/sqrt(length(percentEdited_filterfail(Is)));
    end
    stats(isnan(stats)) = 0;
    varNames = ["Num of Ib cells with edit but failed filters",...
            "Mean reads in Ib cells with edit but failed filters",...
            "Mean Ib editing in Ib cells with edit but failed filters",...
            "Std Dev of Ib editing in Ib cells with edit but failed filters",...
            "Num of Is cells with edit but failed filters",...
            "Mean reads in Is cells with edit but failed filters",...
            "Mean Is editing in Is cells with edit but failed filters",...
            "Std Dev of Is editing in Is cells with edit but failed filters"];
    stats_failed = array2table(stats,'VariableNames',varNames);
    data_processed = [data_processed,stats_failed];
end

function input = addFlybaseID(input,path)

genemap = generateGeneMap(path);

% Compare filterededitssummary locations and record gene name/info
pos = string(input.("Edit Position"));
temp = split(pos,":");
rna.chr = categorical(temp(:,1));
rna.pos = str2double(temp(:,2));
clear temp pos

Gene = strings(size(rna.pos));  %initialize
FlybaseID = strings(size(rna.pos)); %initialize
DNAstrand = zeros(size(rna.pos)); %initialize


for i = 1:size(input,1) % go through each summary position
    index_positionmatch = (genemap.start <= rna.pos(i)) & (rna.pos(i) <= genemap.end);
    index_chrmatch = genemap.chr == rna.chr(i);

    index_gene = index_chrmatch & index_positionmatch;

    if sum(index_gene) == 1 %one gene found at this location
        Gene(i) = genemap{index_gene,"current_symbol"};
        FlybaseID(i) = genemap{index_gene,"primary_FBid"};
        DNAstrand(i) = genemap{index_gene,"dnastrd"};

    else    
        % If more than one gene overlaps at this location, then duplicate
        % the editing position so that each gene is considered
        % independently
        index_multi = find(index_gene);

        for ii = 1:size(index_multi,1)
            
            % On the first pass, record regular info for first gene found
            if ii == 1
                Gene(i) = genemap{index_multi(ii),"current_symbol"};
                FlybaseID(i) = genemap{index_multi(ii),"primary_FBid"};
                DNAstrand(i) = genemap{index_multi(ii),"dnastrd"};
            end

            % On subsequent passes, append new info to bottom of list
            if ii > 1
                input(end+1,:) = input(i,:);
                Gene(end+1) = genemap{index_multi(ii),"current_symbol"};
                FlybaseID(end+1) = genemap{index_multi(ii),"primary_FBid"};
                DNAstrand(end+1) = genemap{index_multi(ii),"dnastrd"}; 
            end
        end
    end

    if mod(i,100)==0; disp(num2str(i)); end
end

input = addvars(input,Gene,'Before','Edit Position');
input = addvars(input,FlybaseID,'After','Gene');
input = addvars(input,DNAstrand,'After','FlybaseID');

clear i Gene FlybaseID DNAstrand gene_dual fb_dual strnd_dual index_gene index_positionmatch index_chrmatch

end

function input = addGeneSummary(input)
    % Download "best gene summaries" from here https://flybase.org/downloads/bulkdata

    % Set up the Import Options and import the data
    opts = delimitedTextImportOptions("NumVariables", 4);
    
    % Specify range and delimiter
    opts.DataLines = [1, Inf];
    opts.Delimiter = "\t";
    
    % Specify column names and types
    opts.VariableNames = ["FBgn_ID", "Var2", "Var3", "Summary"];
    opts.SelectedVariableNames = ["FBgn_ID", "Summary"];
    opts.VariableTypes = ["string", "string", "string", "string"];
    
    % Specify file level properties
    opts.ExtraColumnsRule = "ignore";
    opts.EmptyLineRule = "read";
    
    % Specify variable properties
    opts = setvaropts(opts, ["FBgn_ID", "Var2", "Var3", "Summary"], "WhitespaceRule", "preserve");
    opts = setvaropts(opts, ["FBgn_ID", "Var2", "Var3", "Summary"], "EmptyFieldRule", "auto");
    
    % Import the data
    summaries = readtable("C:\Users\Littleton Lab\Documents\Andres C drive\RNA editing\Flybase genomic data\Best gene summary\best_gene_summary_fb_2023_04.tsv", opts);

    % Clear temporary variables
    clear opts

    % Delete extra rows
    summaries(1:9,:) = [];
    

    % For each RNA edit in the input, find the corresponding summary
    for i = 1:size(input,1)
        index = strcmp(summaries.FBgn_ID,input{i,"FlybaseID"});
        
        GeneSummary(i) = summaries{index,"Summary"};
    end

    % Save summaries into new table variable
    input = addvars(input,GeneSummary','NewVariableNames','GeneSummary','After','PfamDomain');
end

function genemap = generateGeneMap(path)

    % Download from flybase the genes map table
    % https://flybase.org/downloads/bulkdata
    % and import it into matlab, save as a matlab table text columns
    
    
    % Set up the Import Options and import the data
    opts = delimitedTextImportOptions("NumVariables", 6);
    
    % Specify range and delimiter
    opts.DataLines = [1, Inf];
    opts.Delimiter = "\t";
    
    % Specify column names and types
    opts.VariableNames = ["Var1", "current_symbol", "primary_FBid", "Var4", "Var5", "sequence_loc"];
    opts.SelectedVariableNames = ["current_symbol", "primary_FBid", "sequence_loc"];
    opts.VariableTypes = ["string", "string", "string", "string", "string", "string"];
    
    % Specify file level properties
    opts.ExtraColumnsRule = "ignore";
    opts.EmptyLineRule = "read";
    
    % Specify variable properties
    opts = setvaropts(opts, ["Var1", "current_symbol", "primary_FBid", "Var4", "Var5", "sequence_loc"], "WhitespaceRule", "preserve");
    opts = setvaropts(opts, ["Var1", "current_symbol", "primary_FBid", "Var4", "Var5", "sequence_loc"], "EmptyFieldRule", "auto");
    
    % Import the data
    genemap = readtable([path.flybasedata '\Gene map\gene_map_table_fb_2023_04.tsv'], opts);
    
    % Clear temporary variables
    clear opts


    % Remove genes which are not mapped
    seq = [genemap.sequence_loc];
    unmapped_loci = ismissing(seq) | seq == "";
    genemap = genemap(~unmapped_loci,:);
    genemap(1,:) = [];
    
    
    
    %Split sequence_loc column
    temp = split(genemap.sequence_loc,[":","..","(",")"]);
    genemap.chr = categorical(temp(:,1));
    genemap.start = str2double(temp(:,2));
    genemap.end = str2double(temp(:,3));
    genemap.dnastrd = str2double(temp(:,4));

end

function input = checkForNeighbors(input)

    % Sort input by chromosome, and then by position
    input = sortrows(input,["Chromosome" "Position"]);

    % Calculate distance from previous position
    chrs = unique(input.Chromosome);
    distances_accum = [];
    
    for i = 1:size(chrs,1)
        
        % Find indeces for current chromosome
        index = strcmp(input.Chromosome,chrs(i));
        start = find(index,1);
        stop = find(index,1,'last');

        % Calculate distance from each position to next for this chr
        distanceToNext = input{start+1:stop,"Position"} - input{start:stop-1,"Position"};
        distanceToNext = [distanceToNext; nan];

        % Calculate the minimum distance from the next vs the prev position
        distanceFromPrev = [nan; distanceToNext(1:end-1)];
        distances = min(distanceToNext,distanceFromPrev);
        

        if isempty(distances_accum)
            distances_accum = distances;
        else
            distances_accum = [distances_accum; distances];
        end

    end
    
    input = addvars(input,distances_accum,'NewVariableNames','DistFromClosestEdit','After','Position');
end

function input = markCanonical(input)

    % For all edits mapped to minus strand genes, the biological change is
    % the reverse complement of the original base
    index.minusStrand = input.DNAstrand == -1;

    base.refcombined = join(string(input.REF(index.minusStrand)),"");
    base.refcombined_comp = seqcomplement(base.refcombined);

    base.altcombined = join(string(input.ALT(index.minusStrand)),"");
    base.altcombined_comp = seqcomplement(base.altcombined);

    BiologicalBaseChange = string([char(input.REF) char(input.ALT)]);
    BiologicalBaseChange(index.minusStrand) = string([base.refcombined_comp' base.altcombined_comp']);

    input = addvars(input,BiologicalBaseChange,'NewVariableNames','BiologicalBaseChange','After','ALT');

    % Canonical changes are defined as follows:
    index.canonical = ismember(input.BiologicalBaseChange, {'AG'});

    input = addvars(input, index.canonical,'NewVariableNames','Canonical','After','BiologicalBaseChange');
end

function data_summary = coverage(data_summary, data_edits)

    % Load flybase data
    f = waitbar(0,'Calculating read depth at edit sites');
    
    % Create a waitbar
    D = parallel.pool.DataQueue;
    afterEach(D, @nUpdateWaitbar);
    waitbar(0,f,'Calculating read depth at edit sites');
    global progress;
    progress = 0;
    send(D,[progress size(data_summary,1)]);

    % Load read depths (previously calculated in GATK for these locations)
        opts = delimitedTextImportOptions("NumVariables", 4);
        opts.DataLines = [2, Inf];
        opts.Delimiter = ["\t", " ", ":"];
        opts.VariableNames = ["Edit", "Position", "Cell", "Depth"];
        opts.VariableTypes = ["string", "double", "string", "double"];
        opts.ExtraColumnsRule = "ignore";
        opts.EmptyLineRule = "read";
        opts.ConsecutiveDelimitersRule = "join";
        opts = setvaropts(opts, ["Edit", "Cell"], "WhitespaceRule", "preserve");
        opts = setvaropts(opts, ["Edit", "Cell"], "EmptyFieldRule", "auto");

        filedir = "C:\Users\Littleton Lab\Documents\Andres C drive\RNA editing\GATK analysis - Muscle vs neurons\GATK output\Coverage\Read depth at specific locations\";
        batch1 = readtable(append(filedir,"neuron_RNAseq061819_coverage_at_specific_locations.txt"), opts);
        batch2 = readtable(append(filedir,"neuron_RNAseq082119_coverage_at_specific_locations.txt"), opts);
        batch3 = readtable(append(filedir, "neuron_RNAseq091219_coverage_at_specific_locations.txt"), opts);
        muscle = readtable(append(filedir, "muscle_coverage_at_specific_locations.txt"), opts);

        clear opts

        % Combine read depths into single table
        readdepth = [batch1; batch2; batch3; muscle];
        clear batch1 batch2 batch3 muscle

        % Change cellID to standard format
        readdepth.Cell = append("d",extractAfter(readdepth.Cell, "D19-"));

        % Combine Chr and Location
        readdepth.Location = append(readdepth.Edit,":",string(readdepth.Position));

    % Load CellType information

        %Load neuron
        opts = spreadsheetImportOptions("NumVariables", 2);
        opts.Sheet = "Sheet1";
        opts.DataRange = "A2:B207";
        opts.VariableNames = ["CellName", "CellType"];
        opts.VariableTypes = ["string", "string"];
        opts = setvaropts(opts, ["CellName", "CellType"], "WhitespaceRule", "preserve");
        opts = setvaropts(opts, ["CellName", "CellType"], "EmptyFieldRule", "auto");
        
        filedir2 = "C:\Users\Littleton Lab\Documents\Andres C drive\RNA editing\GATK analysis - Muscle vs neurons\Other data files\";
        CellTypeGuide = readtable(append(filedir2, "CellTypeGuide.xlsx"), opts, "UseExcel", false);
        
        clear opts
        CellTypeGuide.CellName = extractBefore(CellTypeGuide.CellName,8);

        % Load muscle
        opts = spreadsheetImportOptions("NumVariables", 2);
        opts.Sheet = "Sheet1";
        opts.DataRange = "A1:B43";
        opts.VariableNames = ["CellName", "CellType"];
        opts.VariableTypes = ["string", "string"];
        opts = setvaropts(opts, ["CellName", "CellType"], "WhitespaceRule", "preserve");
        opts = setvaropts(opts, ["CellName", "CellType"], "EmptyFieldRule", "auto");
        
        muscleIDs = readtable(append(filedir2, "Muscle Cells RNAseq ID numbers.xlsx"), opts, "UseExcel", false);
        CellTypeGuide = [CellTypeGuide; muscleIDs];
        clear opts muscleIDs filedir filedir2


    % Only include read depth for cells we used for RNA sequencing
        Lia = ismember(readdepth.Cell,CellTypeGuide.CellName);
        readdepth = readdepth(Lia,:);

    % Add cell type information to read depth table
        [~,Locb] = ismember(readdepth.Cell,CellTypeGuide.CellName);
        readdepth.CellType = CellTypeGuide.CellType(Locb);
            
    % Find indeces for Ib and Is cells
    idx.Ib = strcmp(readdepth.CellType,"Ib");
    idx.Is = strcmp(readdepth.CellType,"Is");
    idx.M = strcmp(readdepth.CellType,"M");
        

    % Add new columns to data table for read depth values
        Ib = nan(size(data_summary,1),1);
        Is = nan(size(data_summary,1),1);
        M  = nan(size(data_summary,1),1);

        Ib_num = nan(size(data_summary,1),1);
        Is_num = nan(size(data_summary,1),1);
         M_num = nan(size(data_summary,1),1);

    % Slices for parfor loop
        positions = data_summary.("Edit Position");
        readdepthlocation = readdepth.Location;
        dataeditsposition = data_edits.("Edit Position");
        dataeditscellid = data_edits.CellID;

    % For each edit position in data_summary, calculate average Ib and Is
    % read depth, excluding cells that made it past RNA edit filters
        parfor i = 1:size(data_summary,1)   %parfor loop
            currentEdit = positions(i);
        
            % Find indeces of this edit position using logical indexing
            idx_currentPos = readdepthlocation == currentEdit;

            % Find indeces for cells with the edit (both Ib, Is, and M)
            idx_cellswithedit_dataedits = dataeditsposition == currentEdit;
            namesofcellswithedit = dataeditscellid(idx_cellswithedit_dataedits);
            idx_cellswithedit = ismember(readdepth.Cell,namesofcellswithedit);


            % Calculate Ib and Is read depth
            index_currentIbs = idx_currentPos & idx.Ib;
            index_currentIbs = index_currentIbs & ~idx_cellswithedit;

            Ib_num(i) = size(unique(readdepth.Cell(index_currentIbs)),1);
            Ib(i) = mean(readdepth.Depth(index_currentIbs));

            index_currentIss = idx_currentPos & idx.Is;
            index_currentIss = index_currentIss & ~idx_cellswithedit;

            Is(i) = mean(readdepth.Depth(index_currentIss));
            Is_num(i) = size(unique(readdepth.Cell(index_currentIss)),1);

            index_currentMs = idx_currentPos & idx.M;
            index_currentMs = index_currentMs & ~idx_cellswithedit;

            M(i) = mean(readdepth.Depth(index_currentMs));
            M_num(i) = size(unique(readdepth.Cell(index_currentMs)),1);
         
            % Update waitbar
            send(D, [i,size(data_summary,1)]);
            %disp(append("Coverage for unedited cell. Calculating edit #: ",num2str(i)))
        end

    % Add read depth info to data_summary
    data_summary = addvars(data_summary,Ib_num, Ib,...
        'After','Num of Ib cells with edit',...
        'NewVariableNames',{'Num of Ib cells without edit','Mean reads in Ib cells without edit'});
    data_summary = addvars(data_summary,Is_num, Is,...
        'After','Num of Is cells with Edit',...
        'NewVariableNames',{'Num of Is cells without edit','Mean reads in Is cells without edit'});
    data_summary = addvars(data_summary,M_num, M,...
        'After','Num of M samples with edit',...
        'NewVariableNames',{'Num of M cells without edit','Mean reads in M cells without edit'});
    
    close(f);

end
```
