## Supplementary material for "A stochastic RNA editing process targets a select number of sites in individual *Drosophila* glutamatergic motoneurons": MATLAB code: Step4_Graphs_and_tables.m

```
% A stochastic RNA editing process targets a limited number of sites in individual Drosophila glutamatergic motoneurons
% Andrés B. Crane, Suresh K. Jetti, J. Troy Littleton
% The Picower Institute for Learning and Memory, Department of Brain and Cognitive Sciences, Department of Biology, Massachusetts Institute of Technology, Cambridge, MA 02139
% Correspondence and requests for materials should be addressed to J.T.L..


%% Set up workspace and environment
clear
clc

% Set up main directory for loading and saving files
path.main = 'C:\Users\Littleton Lab\Documents\Andres C drive\RNA editing\GATK analysis - Muscle vs neurons\';

% Only graph canonical sites
canonical = false;

if canonical == true
    path.results = [path.main '\Results\Canonical'];
else
    path.results = [path.main '\Results\Noncanonical'];
end

path.graphs = [path.results '\Graphs'];
path.tables = [path.results '\Tables'];

load([path.main '\Results\Matlab files.mat'])
biochange = data_summary.BiologicalBaseChange;
data_summary = data_summary(data_summary.Canonical==canonical,:);
data_edits = data_edits(data_edits.Canonical==canonical,:);

% Only graph neuron edits
indexN = contains(data_summary.("Cell types with edit"),"N");
sum(data_summary.("Cell types with edit") == "M")
data_summary = data_summary(indexN,:);


% set default figure values
markersize = 100;
myfontsize = 9;
myfont = 'Arial';
set(groot,'defaultFigureColor','w')
set(groot,'defaultAxesfontname',myfont)
set(groot, 'DefaultAxesFontSize', myfontsize);
set(groot, 'defaultFigureUnit', 'Inches');


color_Ib = [56 126 142]./255;
color_Is = [220 145 57]./255;


%% Statistics about data
% Mutation type
groupsummary(data_summary,'MutationType')

%Number of unique genes
size(unique(data_summary.Gene),1)

% Novel sites
groupsummary(data_summary,'PreviouslyKnownRNAeditingSite')

% Gene part distribution
groupsummary(data_summary,'GenePart')

% Significant edits
groupsummary(data_summary,'AminoAcidGroupChange')

% Penetrance of edits
max(data_summary.("Num of Ib cells + num of Is cells"))
mean(data_summary.("Num of Ib cells + num of Is cells"))
%% Num of cells with edit scatter plot

fig = figure(2);
s = scatter(data_summary.("Num of Ib cells with edit")./105,...
            data_summary.("Num of Is cells with Edit")./101,[],...
            data_summary.("Mean Ib editing - mean Is editing"),'.');
r = corrcoef(data_summary.("Num of Ib cells with edit")./105,...
            data_summary.("Num of Is cells with Edit")./101)
%title("Fraction of cells colored by mean editing difference")
xlabel("Fraction of Ib cells with edit");ylabel("Fraction of Is cells with edit")
%xlim([0 105]);ylim([0 105]);
axis square
colormap jet
cb = colorbar;
cb.Label.String = "Mean Ib editing - mean Is editing";
s.SizeData = markersize;

%orient(figure(2),'landscape')

set(gca,'box','off');
fig.Position = [0 1 2.5 2];
fontsize(fig,myfontsize,"Points")
fontname(fig,myfont);
exportgraphics(fig,[path.graphs '\NumOfCellsWithEditScatter.pdf'])

%% percent editing scatter plot
fig = figure(3);

s = scatter(data_summary.("Mean Ib editing"),data_summary.("Mean Is editing"),'b.');
r = corrcoef(data_summary.("Mean Ib editing"),data_summary.("Mean Is editing"))

%title("Mean editing")
xlabel("Mean Ib editing");ylabel("Mean Is editing")
axis square
lsline
s.SizeData = markersize;
set(gca,'box','off');
fig.Position = [0 1 2.5 2];
fontsize(fig,myfontsize,"Points")
fontname(fig,myfont);
exportgraphics(fig,[path.graphs '\PercentEditingScatter.pdf'])
%% Genes with the most edit sites
fig = figure(4);


% Overall
G = groupsummary(data_summary,'Gene');
G = sortrows(G,'GroupCount','descend');

range = (1:20);
x = categorical(G.Gene(range));
x = reordercats(x,string(x));
bar(x,G.GroupCount(range),'FaceAlpha',0.6);
%title('Genes with the most edit sites')
ylabel('Number of edits')
xtickangle(45)
fontsize(fig,myfontsize,"Points")
fontname(fig,myfont);
set(gcf,'position',[0,2,4,2.95])
set(gca,'box','off')
exportgraphics(fig,[path.graphs '\Genes with the most edit sites.pdf'])
%% Most penetrant edits

% Figure 
fig = figure(5);
penetrant = sortrows(data_summary,'Num of Ib cells + num of Is cells','descend');
histogram(penetrant.("Num of Ib cells + num of Is cells"))
xlabel('Number of cells an edit is in');ylabel('Number of edits')
%title('Edit penetrance')
fontsize(fig,myfontsize,"Points")
fontname(fig,myfont);
fig.Position = [0 2 3 2.5];
set(gca,'box','off')
exportgraphics(fig,[path.graphs '\Edit penetrance.pdf'])
%mean(penetrant.("Num of Ib cells + num of Is cells"))

% Figure
range = (1:20);
vars = ["Gene","Edit Position","GenePart","Num of Ib cells with edit", "Num of Is cells with Edit"];
penetrant = penetrant(range,vars);
penetrant.("% of Ib cells with edit") = penetrant.("Num of Ib cells with edit")./105;
penetrant.("% of Is cells with edit") = penetrant.("Num of Is cells with Edit")./101;

writetable(penetrant,[path.tables '\Edit penetrance.xlsx'],'WriteMode','overwrite')

fig = figure();
x2 = penetrant.GenePart;
x2 = replace(x2,"FivePrimeUTR","5'UTR");
x2 = replace(x2,"ThreePrimeUTR","3'UTR");
x = strcat(penetrant.Gene,"-",x2,"-",penetrant.("Edit Position"));
x = categorical(x,x);

y = 100*[penetrant.("% of Ib cells with edit"),penetrant.("% of Is cells with edit")];
b = bar(categorical(x),y); %,'FaceColor',color_Ib)

set(gca,'box','off')
b(1).FaceColor = color_Ib;
b(2).FaceColor = color_Is;
L = legend('Ib','Is');
fig.Position = [0 2 8 4];
ylabel('Percent of cells with edit')
fontsize(fig,myfontsize,"Points")
fontname(fig,myfont);

exportgraphics(fig,[path.graphs '\Edit penetrance - top edits.pdf'])
%% Ib and Is vs editing percentage
fig = figure(6);
nbins = 30;
h = histogram(data_summary.("Mean Ib editing"),nbins);
h.FaceColor = color_Ib;
hold on
h = histogram(data_summary.("Mean Is editing"),nbins);
h.FaceColor = color_Is;
hold off

%title('Edit percentage in Ib vs Is')
xlabel('Mean fraction of reads edited')
ylabel('Number of edit sites')
L = legend('Ib','Is');
L.Location = 'northwest';

Ibmean = mean(data_summary.("Mean Ib editing"));
Ismean = mean(data_summary.("Mean Is editing"));
[~,p] = ttest2(data_summary.("Mean Ib editing"),data_summary.("Mean Is editing"));

%text(0.7,0.8,['Ib mean = ' num2str(Ibmean)],'units','normalized')
%text(0.7,0.75,['Is mean = ' num2str(Ismean)],'units','normalized')
%text(0.7,0.7,['pval ttest = ' num2str(p)],'units','normalized')

set(gca,'box','off');
fig.Position = [0 1 2.5 2];
fontsize(fig,myfontsize,"Points")
fontname(fig,myfont);

exportgraphics(fig,[path.graphs '\EditPercentageIbvsIsHistogram.pdf'])
%% Edits most likely to have coding impact
fig = figure(7);
impact = data_summary(data_summary.AminoAcidGroupChange=="Yes",:);
writetable(impact,[path.tables '\ImpactfulCDSEdits.xlsx'],'WriteMode','overwrite');

Ib_v_Is_numcells = impact.("Num of Ib cells with edit")-impact.("Num of Is cells with Edit");
Ib_v_Is_editing = impact.("Mean Ib editing")-impact.("Mean Is editing");
combined = table;
combined.Ib_v_Is_numcells = Ib_v_Is_numcells;
combined.Ib_v_Is_editing = Ib_v_Is_editing;
combined.Gene = impact.Gene;


s = scatter(combined.Ib_v_Is_numcells,combined.Ib_v_Is_editing,'.');
title('Edits that change the AA group')
xlabel('Num of Ib cells - num of Is cells')
ylabel('Editing in Ib - editing in Is')
s.SizeData = markersize;

range = 1:5;
combined = sortrows(combined,'Ib_v_Is_numcells','descend');
text(combined.Ib_v_Is_numcells(range)+1,combined.Ib_v_Is_editing(range),combined.Gene(range))

combined = sortrows(combined,'Ib_v_Is_editing','descend');
text(combined.Ib_v_Is_numcells(range)+1,combined.Ib_v_Is_editing(range),combined.Gene(range))

combined = sortrows(combined,'Ib_v_Is_editing','ascend');
text(combined.Ib_v_Is_numcells(range)+1,combined.Ib_v_Is_editing(range),combined.Gene(range))


exportgraphics(fig,[path.graphs '\AAgroupChangeCDSedits.pdf'])
%% Data validation - how many previously known edit sites pop up
prevknown = data_summary.PreviouslyKnownRNAeditingSite=="Yes";
sum(prevknown)


%% Does one cell have high editing across many sites?

% Find IDs of cells that have edits, and sort so Ibs and Iss are together
cellnames = unique(string([data_edits.CellID data_edits.CellType]),'rows');
cellnames = sortrows(cellnames,2);
cellnames = num2cell(cellnames);

% For each cell, extract the fraction of reads edited at each site
numCells = size(cellnames,1);
for row = 1:numCells
    thisCellsEditsIndeces = strcmp(data_edits.CellID,cellnames{row});
    cellnames(row,3) = {data_edits.FractionOfReadsEdited(thisCellsEditsIndeces)};
end

% Find the average fraction of reads edited, and sort by this within Ib and
% Is
cellnames(:,4) = num2cell(cellfun(@mean,cellnames(:,3)));

indeces_Ib = strcmp([cellnames{:,2}],"Ib");
indeces_Is = strcmp([cellnames{:,2}],"Is");
indeces_M = strcmp([cellnames{:,2}],"M");

cellnames(indeces_Ib,:) = sortrows(cellnames(indeces_Ib,:),4);
cellnames(indeces_Is,:) = sortrows(cellnames(indeces_Is,:),4);
cellnames(indeces_M,:) = sortrows(cellnames(indeces_M,:),4);

% Plot
fig = figure(9);
xyaccum=[];
for row = 1:numCells
    yvals = cellnames{row,3};
    xval = ones(size(yvals))*row;
    xyaccum = [xyaccum; xval yvals];

    %swarmchart(xval,yvals,'b.')
    %hold on
end

% Plot a scatter of each point, where the color of the point is set by the
% density of nearby points
x = xyaccum(:,1);
y = xyaccum(:,2);
c = ksdensity(xyaccum,xyaccum);
%c = ksdensity([x,y], [x,y]);
s = scatter(x, y, [], c,'.');
%s.SizeData = 50;
hold on

% Add red line for average
celltypediv = unique([cellnames{:,2}]);
plot(1:sum(indeces_Ib),[cellnames{indeces_Ib,4}],'r','LineWidth',2)
plot(sum(indeces_Ib)+1:(sum(indeces_Is)+sum(indeces_Ib)),[cellnames{indeces_Is,4}],'r','LineWidth',2)
%plot(sum(indeces_Ib)+1:(sum(indeces_Is)+sum(indeces_Ib)),[cellnames{indeces_Is,4}],'r','LineWidth',2)


% Figure formatting
%title("Does one cell have high editing across many sites?")
xlabel("Single cells (1-105 Ib, 106-206 Is)")
ylabel(["Fraction of reads edited " "(each dot is one edit site)"])
%xlim([0 249])  %use this to also plot muscles
xlim([0 206])
%L(1) = plot(nan, nan, 'b.');
%L(2) = plot(nan, nan, 'r.','LineWidth',2);
%legend(L, {'Single edits', 'Average editing of cell'})
L = legend('Single edits', 'Average editing of cell');
L.Location = 'southeast';
L.ItemTokenSize(1) = 10;
colormap(jet)
hold off


orient(figure(9),'landscape')

cb = colorbar;
cb.Label.String = "Point density (kernel density estimation)";
fig.Position = [0 1 8.5 3.5];
fontsize(fig,myfontsize,"Points")
fontname(fig,myfont);

exportgraphics(figure(9),[path.graphs '\AverageEditingAcrossAllCells.pdf'])

%% Pie chart for GenePart
fig = figure(10);
data = data_summary.GenePart;
data = replace(data,"FivePrimeUTR","5'UTR");
data = replace(data,"ThreePrimeUTR","3'UTR");
p = pie(categorical(data));
%p = pie(categorical(data_summary.GenePart));
%title("Edit % in mRNA region")

% New formatting
fig.Position = [0 0 2.8 1.8];
fontsize(fig,myfontsize,"Points")
fontname(fig,myfont);
p(1).FaceColor = [39/255, 170/255, 225/255];    % 3'UTR
p(5).FaceColor = [0/255, 166/255, 81/255];      % CDS
p(3).FaceColor = [251/255, 176/255, 64/255];    % 5'UTR

%old formatting
%set(findobj(p,'type','text'),'fontsize',25, 'fontname','arial')
%set(gca,'FontSize',25)
%set(gcf,'position',[70,70,1000,500])

exportgraphics(fig,[path.graphs '\GenePart.pdf'])

%% Pie chart for CDS sub-categories (type of mutation)

% Figure 11 - Distribution of CDS edits
fig = figure(11);
idx = strcmp(data_summary.MutationType,"-");
MutationType = data_summary.MutationType(~idx);
p = pie(categorical(MutationType));
%title("Distribution of CDS edits")
%set(findobj(p,'type','text'),'fontsize',8, 'fontname','arial')
%set(gca,'FontSize',25)


% Formatting for fig 11
set(gcf,'position',[0,0,3.1,1.7])
fontsize(fig,myfontsize,"Points")
fontname(fig,myfont);
p(1).FaceColor = [41/255, 171/255, 104/255];
p(3).FaceColor = [140/255, 222/255, 180/255];
exportgraphics(figure(11),[path.graphs '\Distribution of CDS edits.pdf'])


% Figure 12 - Distribution of missense edits
fig = figure(12);
idx2 = strcmp(data_summary.MutationType,"Missense");
AAgroupchange = data_summary.AminoAcidGroupChange(idx2);
p = pie(categorical(AAgroupchange));

% Formatting for fig 12
fontsize(fig,myfontsize,"Points")
fontname(fig,myfont);
set(gcf,'position',[0,0,2.8,1.8])
%title(["Distribution of missense edits" "affecting amino acid group type"])
p(3).FaceColor = [255/255, 129/255, 94/255];
p(1).FaceColor = [255/255, 189/255, 172/255];
exportgraphics(figure(12),[path.graphs '\Dist of missense edits.pdf'])
%% Do all edits happen near beginning of 3' or 5' UTR?
fig = figure(13);
tiledlayout(2,1);
nexttile;
utr3indeces = data_summary.GenePart == "ThreePrimeUTR";
x = data_summary{utr3indeces,"PosInGenePart"};
histogram(x,1:100:max(x),'Normalization','probability');
%[N,edges] = histcounts(data_summary{utr3indeces,"PosInGenePart"},1:100:5000,'Normalization','probability');
title('Position of edits in 3'' UTR')
xlabel('Nucleotide Position');
ylabel('Probability')

nexttile;
x = data_summary{utr3indeces,"PosInGenePartFrac"};
histogram(x,20,'Normalization','probability');
xlabel('Position along 3''UTR (fraction)');
ylabel('Probability')

exportgraphics(figure(13),[path.graphs '\Position of edits in 3'' UTR.pdf'])


fig = figure(14);
tiledlayout(2,1);
nexttile;
utr5indeces = data_summary.GenePart == "FivePrimeUTR";
x = data_summary{utr5indeces,"PosInGenePart"};
histogram(x,1:100:max(x),'Normalization','probability');
%[N,edges] = histcounts(data_summary{utr3indeces,"PosInGenePart"},1:100:5000,'Normalization','probability');
title('Position of edits in 5'' UTR')
xlabel('Nucleotide Position');
ylabel('Probability')

nexttile;
x = data_summary{utr5indeces,"PosInGenePartFrac"};
histogram(x,20,'Normalization','probability');
xlabel('Position along 5''UTR (fraction)');
ylabel('Probability')

exportgraphics(figure(14),[path.graphs '\Position of edits in 5'' UTR.pdf'])
%% Edit probablity vs gene expression level
fig = figure(15);
s = scatter(data_summary.("Gene TPM in Ib"),data_summary.("Mean Ib editing"),'.');
R_Ib = corrcoef(data_summary.("Gene TPM in Ib"),data_summary.("Mean Ib editing"));
%s.SizeData = markersize;
s(1).CData = color_Ib;
s(1).MarkerFaceColor= color_Ib;
hold on
s = scatter(data_summary.("Gene TPM in Is"),data_summary.("Mean Is editing"),'.');
R_Is = corrcoef(data_summary.("Gene TPM in Is"),data_summary.("Mean Is editing"));
%s.SizeData = markersize;
s(1).CData = color_Is;
s(1).MarkerFaceColor= color_Is;
hold off
h = lsline;
set(h(1),'color',color_Ib)
set(h(2),'color',color_Is)

% Figure formatting
L = legend('Ib','Is');
%title('Does edit probability depend on gene expression level?')
xlabel('Gene log_{2}TPM')
ylabel('Fraction of reads edited')
L.ItemTokenSize(1) = 10;

fig.Position = [0 1 2.7 2];
if canonical == false
    fig.Position = [0 1 2.7 2.35];
end
fontsize(fig,myfontsize,"Points")
fontname(fig,myfont);
exportgraphics(figure(15),[path.graphs '\Edit probablity vs gene expression level.pdf'])

%% Edits in Ib that are not in Is (and vice versa)
fig = figure(16);
only_Ib = data_summary(data_summary.('Mean Is editing')==0,:);
only_Is = data_summary(data_summary.('Mean Ib editing')==0,:);

s = scatter(only_Ib.("Mean Ib editing"),only_Ib.("Mean Is editing"),'.');
s.MarkerFaceColor = color_Ib;
s.SizeData = markersize;
h = text(only_Ib.("Mean Ib editing"),only_Ib.("Mean Is editing")+0.05,only_Ib.Gene);
set(h,'Rotation',60);
%text(combined.Ib_v_Is_numcells(range)+1,combined.Ib_v_Is_editing(range),combined.Gene(range));
hold on
s = scatter(only_Is.("Mean Ib editing"),only_Is.("Mean Is editing"),'.');
s.MarkerFaceColor = color_Is;
s.SizeData = markersize;
text(only_Is.("Mean Ib editing")+0.05,only_Is.("Mean Is editing"),only_Is.Gene);
hold off
title("Mean editing only in Ib or Is")
xlabel("Mean Ib editing");ylabel("Mean Is editing")
axis square
l = legend('Only edited in Ib','Only edited in Is');

exportgraphics(figure(16),[path.graphs '\Mean editing only in Ib or Is.pdf'])

%% Editing vs ADAR expression
% fig = figure(17);
% 
% % Set the color of each point by the TPM expression
% c = data_edits.("ADAR TPM");
% 
% 
% s = scatter(2.^data_edits.("ADAR TPM"),data_edits.FractionOfReadsEdited,[],c,'.');
% s.SizeData = markersize;
% title('{\it adar} expression level vs RNA editing')
% ylabel('Fraction of Reads Edited');
% xlabel('{\it adar} expression level (TPMs)')
% lsline
% exportgraphics(figure(17),[path.graphs '\ADAR expression level vs RNA editing.pdf'])

%% Editing vs ADAR expression with color
fig = figure(17);


% Find mean TPM expression for each
data_edits.("Gene TPM mean") = mean([data_edits.("Gene TPM in Ib"),data_edits.("Gene TPM in Is")],2);

% Set the color of each point by the TPM expression
red = [1 0 0];
blue = [132 203 255]./255;


fraction = 0.7;
cdata = data_edits{:,"Gene TPM mean"};

indeces = data_edits.("Gene TPM mean")>fraction*max(cdata);

mymap = repmat(blue,[100,1]);
indeces_cmap = 100*fraction:100;
mymap(indeces_cmap,:) = repmat(red,[size(indeces_cmap,2),1]);
colormap(mymap);

clim(gca,[0 max(cdata)])

% Plot scatter
s = scatter(data_edits.("ADAR TPM"),data_edits.FractionOfReadsEdited,[],blue,'.');
h1 = lsline;
h1.Color = 'k';
h1.LineWidth = 1;
hold on
s2 = scatter(data_edits{indeces,"ADAR TPM"},data_edits{indeces,"FractionOfReadsEdited"},[],red,'.');
hold off
xlim([0 7])

% Figure formatting
%s.SizeData = markersize;
%title('{\it adar} expression level vs RNA editing')
ylabel('Fraction of reads edited');
xlabel('{\it adar} expression level (log_{2}TPMs)')
cb = colorbar;
cb.Label.String = ["Normalized" "gene expression level"];
fig.Position = [0 1 2.9 2];
fontsize(fig,myfontsize,"Points")
fontname(fig,myfont);

exportgraphics(figure(17),[path.graphs '\ADAR expression level vs RNA editing.pdf'])

%% Cell number vs gene editing level
fig = figure(18);
s = scatter(data_summary.("Num of Ib cells with edit")./105,data_summary.("Mean Ib editing"),'.');
s.SizeData = markersize;
s.CData = color_Ib;
hold on
s = scatter(data_summary.("Num of Is cells with Edit")./101,data_summary.("Mean Is editing"),'.');
s.SizeData = markersize;
s.CData = color_Is;
hold off
h = lsline;
set(h(1),'color',color_Ib)
set(h(2),'color',color_Is)

% Figure formatting
L = legend('Ib','Is');
L.ItemTokenSize(1) = 10;
%title('Gene editing level vs fraction of cells with edit')
xlabel('Fraction of cells with edit')
ylabel('Fraction of reads edited')
fig.Position = [0 1 2.7 2.3];
fontsize(fig,myfontsize,"Points")
fontname(fig,myfont);
exportgraphics(figure(18),[path.graphs '\Gene editing level vs fraction of cells with edit.pdf'])

%% Avg editing in CDS, 3' and 5' UTR
fig = figure(19);
data = data_summary.GenePart;
data = replace(data,"FivePrimeUTR","5'UTR");
data = replace(data,"ThreePrimeUTR","3'UTR");
x = categorical(data);
x = reordercats(x,["5'UTR","CDS","3'UTR"]);


%x = categorical(data_summary.GenePart,["CDS" "FivePrimeUTR" "ThreePrimeUTR"]);
y = data_summary.("Mean Ib editing");
s = swarmchart(x,y,'.');
%s.SizeData = markersize;
s.CData = color_Ib;

hold on
y = data_summary.("Mean Is editing");
s = swarmchart(x,y,'.');
%s.SizeData = markersize;
s.CData = color_Is;
hold off


%[~, objh] = legend('Ib','Is');
%objhl = findobj(objh, 'type', 'line');
%set(objhl, 'Markersize', markersize);
%title('Mean editing in gene region')
ylabel('Fraction of reads edited')
fontsize(fig,myfontsize,"Points")
fontname(fig,myfont);
fig.Position = [0 0 2.5 2];


% ----------------------------------
% Calculate ANOVA p-value between 5'UTR, 3'UTR, and CDS editing

% Make indeces for each gene part
Fpr_i = data_summary.("GenePart") == "FivePrimeUTR";
Tpr_i = data_summary.("GenePart") == "ThreePrimeUTR";
CDS_i = data_summary.("GenePart") == "CDS";

% Make single matrix that combines Ib and Is editing for each gene part
Fpr = [data_summary{Fpr_i,"Mean Ib editing"} ; data_summary{Fpr_i,"Mean Is editing"}];
Tpr = [data_summary{Tpr_i,"Mean Ib editing"} ; data_summary{Tpr_i,"Mean Is editing"}];
CDS = [data_summary{CDS_i,"Mean Ib editing"} ; data_summary{CDS_i,"Mean Is editing"}];

% Create combined matrix with categorical factors
comb = [Fpr; Tpr; CDS];
cats = [repmat("5'UTR",size(Fpr,1),1); ...
        repmat("3'UTR",size(Tpr,1),1); ...
        repmat("CDS",size(CDS,1),1)];

% Check if data is normally distributed
clear h p
[h(3),p(3)] = kstest(CDS);
[h(2),p(2)] = kstest(Tpr);
[h(1),p(1)] = kstest(Fpr);

% Run statistical test between the three groups
aov = anova(cats,comb);
groupmeans(aov)
multcompare(aov)


% Add mean line for each group
hold on
w = 0.2;    %horizontal width of line
plot([1-w 1+w],[mean(Fpr) mean(Fpr)],'k')
plot([2-w 2+w],[mean(CDS) mean(CDS)],'k')
plot([3-w 3+w],[mean(Tpr) mean(Tpr)],'k')
hold off

%legend('Ib','Is','Location','southeast')
L = legend('Ib','Is','Position', [0.83 0.20 0.01 0.1]);
L.ItemTokenSize(1) = 10;

exportgraphics(figure(19),[path.graphs '\Mean editing in gene parts.pdf'])

%% ADAR expression in Ib and Is

% Extract ADAR transcripts per million for each single sequenced neuron
if ~exist('tpmdata_adar','var')
path.tpms = 'C:\Users\Littleton Lab\Documents\Andres C drive\RNA editing\RNA editing code (compares RNAseq to DNAseq of parent flies)';
tpmdata_adar = readmatrix([path.tpms '\Jetti et al - Supp Table 1.xlsx'],'Sheet','1a--TPMs','Range','A1822:HJ1822');
tpmdata_adar_Ib = tpmdata_adar(12:116);
tpmdata_adar_Is = tpmdata_adar(118:218);
end

% Figure 20 - ADAR expression in Ib and Is cells sequenced
fig = figure(20);
swarmchart(categorical(repmat("Ib",105,1)),tpmdata_adar_Ib,markersize,color_Ib,'.');
hold on
swarmchart(categorical(repmat("Is",101,1)),tpmdata_adar_Is,markersize,color_Is,'.');
hold off

% Plot horizontal line for mean of each Ib and Is
y1 = mean(tpmdata_adar_Ib);
y2 = mean(tpmdata_adar_Is);
hold on
plot([.8,1.2],[y1 y1],'k')
plot([1.8,2.2],[y2 y2],'k')
hold off

% Calculate p value using t test (but use the one from Jetti et al)
%[~,p] = ttest2(tpmdata_adar_Ib,tpmdata_adar_Is);
%disp(['P-value between ADAR expression of Ib and Is is ' num2str(p)])
text(1.4,7,5,"ns")

% Figure formatting
ylabel('ADAR mRNA level (log_{2}TPM)')
%title('{\it adar} expression')
fontsize(fig,myfontsize,"Points")
fontname(fig,myfont);
fig.Position = [0 0 2.3 1.8];
markersize = 100;
set(gcf, 'Renderer', 'Painters');
exportgraphics(figure(20),[path.graphs '\adar expression.pdf'])

%% Frequency of Amino Acid changes in missense edits
figure(21)  
index = data_summary.MutationType == 'Missense';
change = [data_summary{index,"AminoAcid_Original"},...
          repmat(char(0x2192),sum(index),1),...
          data_summary{index,"AminoAcid_Edited"}];
h = histogram(sort(categorical(cellstr(change))));
h.DisplayOrder = 'descend';
xlabel('Edit-induced amino acid change')
ylabel('Frequency')
%title('Frequency of specific missense edits')


u = unique(data_summary(index,"FlybaseID"));
hold on
%text(h.NumDisplayBins-8,28,['Number of genes: ' num2str(size(u,1))])
hold off

set(gcf,'position',[0,0,10,1.5])
orient(figure(21),'landscape')
set(gca,'FontSize',myfontsize,'box','off')

exportgraphics(figure(21),[path.graphs '\Frequency of specific missense edits.pdf'])

%% Top edits by difference in number of cells edits

numcells = sortrows(data_summary,'Num of Ib cells - num of Is cells','descend');

vars = ["Gene","Edit Position","GenePart","Num of Ib cells with edit", "Num of Is cells with Edit",...
    "Gene TPM in Ib","Gene TPM in Is","Gene TPM adj pval"];
numcells = numcells([1:20 (end-20):end],vars);

writetable(numcells,[path.tables '\Top edits by difference in number of cells with edit.xlsx'],'WriteMode','overwrite')

%% Top Is edits by editing amount
sorted = sortrows(data_summary,'Mean Ib editing - mean Is editing');
sorted(sorted.("Num of Ib cells with edit") == 0,:) = [];
sorted(sorted.("Num of Is cells with Edit") == 0,:) = [];
numEditsToPlot = 10;
fig = figure(22);
fig = boxswarmchart(fig,sorted,data_edits,numEditsToPlot,color_Ib,color_Is,markersize);
title('Ib and Is edits where Is has much higher editing')
exportgraphics(figure(22),[path.graphs '\Ib and Is edits where Is has much higher editing.pdf'])
%% Top Is edits with no Ib edits
sorted = sortrows(data_summary,'Mean Ib editing - mean Is editing');
sorted = sorted(sorted.("Num of Ib cells with edit") == 0,:);
numEditsToPlot = 9;
fig = figure(23);
fig = boxswarmchart(fig,sorted,data_edits,numEditsToPlot,color_Ib,color_Is,markersize);
title('Top Is edits with no Ib editing')
exportgraphics(figure(23),[path.graphs '\Top Is edits with no Ib editing.pdf'])
%% Top Is edits by editing amount
sorted = sortrows(data_summary,'Mean Ib editing - mean Is editing','descend');
sorted(sorted.("Num of Ib cells with edit") == 0,:) = [];
sorted(sorted.("Num of Is cells with Edit") == 0,:) = [];
numEditsToPlot = 10;
fig = figure(24);
fig = boxswarmchart(fig,sorted,data_edits,numEditsToPlot,color_Ib,color_Is,markersize);
title('Ib and Is edits where Ib has much higher editing')
exportgraphics(figure(24),[path.graphs '\Ib and Is edits where Ib has much higher editing.pdf'])
%% Top Ib edits with no Is edits
sorted = sortrows(data_summary,'Mean Ib editing - mean Is editing');
sorted = sorted(sorted.("Num of Is cells with Edit") == 0,:);
numEditsToPlot = 9;
fig = figure(25);
fig = boxswarmchart(fig,sorted,data_edits,numEditsToPlot,color_Ib,color_Is,markersize);
title('Top Ib edits with no Is editing')
exportgraphics(figure(25),[path.graphs '\Top Ib edits with no Is editing.pdf'])
%% Top Ib CDS edits
index = data_summary.GenePart == "CDS";
sorted = sortrows(data_summary(index,:),"Mean Ib editing",'descend');
numEditsToPlot = 10;
fig = figure(26);
fig = boxswarmchart(fig,sorted,data_edits,numEditsToPlot,color_Ib,color_Is,markersize);
title('Top Ib CDS edits')
exportgraphics(figure(26),[path.graphs '\Top Ib CDS edits.pdf'])
%% Top Is CDS edits
index = data_summary.GenePart == "CDS";
sorted = sortrows(data_summary(index,:),"Mean Is editing",'descend');
numEditsToPlot = 10;
fig = figure(27);
fig = boxswarmchart(fig,sorted,data_edits,numEditsToPlot,color_Ib,color_Is,markersize);
title('Top Is CDS edits')
exportgraphics(figure(27),[path.graphs '\Top Is CDS edits.pdf'])
%% Genes with similar TPM expression but diff editing
Nindex = (data_summary.("Num of Ib cells with edit") > 9)...
       | (data_summary.("Num of Is cells with Edit") > 9);

Nonly = data_summary(Nindex,:);

TPMdiff = 2.^data_summary.("Gene TPM in Ib") - 2.^data_summary.("Gene TPM in Is");
data_summary.TPMdiff = TPMdiff;

fig = figure(28);
scatter(data_summary.TPMdiff,data_summary.("Mean Ib editing - mean Is editing"),'.')
%title('Genes with similar TPM expression but diff editing')
xlabel(["Difference in TPMs" "(Ib TPM - Is TPM)" "Is<-------Higher in------->Ib"])
ylabel(["Difference in editing" "(Ib editing - Is editing)" "Is<-------Higher in------->Ib" ])

index = data_summary.("P-val of Ib vs Is editing")<0.05 & ...
        data_summary.("Gene TPM adj pval") > 0.05;
hold on
scatter(data_summary{index, "TPMdiff"},data_summary{index, "Mean Ib editing - mean Is editing"},'r.');
hold off

L = legend('All edit sites','Edit sites with similar TPMs but different editing','Location','southoutside');
L.ItemTokenSize(1) = 10;
%set(gcf,'position',[1800 647 600 550])
set(gca,'box','off');
fig.Position = [0 1 4 3.5];
fontsize(fig,myfontsize,"Points")
fontname(fig,myfont);
exportgraphics(figure(28),[path.graphs '\Genes with similar TPM expression but diff editing.pdf'])

writetable(data_summary(index,:),...
           [path.tables '\Supplemental Table 6 - Differences between Ib and Is.xlsx'],...
           'WriteMode','overwritesheet')


%% Plot histogram of edits

% for i = 1:size(Edited_accum,1)
%     x(i,:) = sum(cat(1,Edited_accum{i,:}),"omitnan");
% end
% categories = ["AC" "AG" "AT"...
%               "CA" "CG" "CT"...
%               "GA" "GC" "GT"...
%               "TA" "TC" "TG"];
% titles = ["All DNA/RNA mismatches",...
%           "Filtered by min reads",...
%           "Filtered by SNPs",...
%           "Filtered by parent lines same DNA",...
%           "Filtered by min edit percentage",...
%           "Filtered by min cell number",...
%           "Filtered by AG/TC only"];
% fig = figure(28);
% fig.Position = [1000 175 560 1163];
% tiledlayout(size(x,1),1)
% for i = 1:size(x,1)
%     nexttile
%     histogram('Categories',categories,'BinCounts',x(i,:))
%     title(titles(i))
%     hold on
% end


%% Closest edit

% VERSION 1 ----------------------------------------------------------

% fig = figure(32);
% t = tiledlayout(2,1,'TileSpacing','Compact');
% 
% nexttile
% histogram(data_summary.DistFromClosestEdit,[0:1:100 Inf])
% ylim([22 400])
% h = gca;
% h.XAxis.Visible = 'off';
% 
% nexttile
% histogram(data_summary.DistFromClosestEdit,[0:1:100 Inf])
% ylim([0 22])
% 
% %title(t,'Distance between edits')
% xlabel(t,'Distance from closest edit (bp)')
% ylabel(t,'Number of occurences')
% fontsize(fig,myfontsize,"Points")
% fontname(fig,myfont);
% fig.Position = [0 2 3.5 2];
% 
% exportgraphics(figure(32),[path.graphs '\Distance between closest edit sites.pdf'])

% To calculate number of sites 1-10 nucleotodes away
% idx = data_summary.DistFromClosestEdit <100;
% t = data_summary.DistFromClosestEdit(idx);
% sum(t<10)/size(t,1)

% Version 2 same figure --------------------------------------------------
fig = figure(32);

yyaxis left
histogram(data_summary.DistFromClosestEdit,[0:1:100 Inf])
%ylim([0 120])

%title(t,'Distance between edits')
xlabel('Distance from closest edit (bp)')
ylabel('Number of occurences')
text(105,-13,"+")


yyaxis right
[f,x] = ecdf(data_summary.DistFromClosestEdit);
plot(x,f)
xlim([0 103])
ylabel('Empirical cumulative density')

fontsize(fig,myfontsize,"Points")
fontname(fig,myfont);
fig.Position = [0 2 3.5 2];
set(gca,'box','off')
exportgraphics(figure(32),[path.graphs '\Distance between closest edit sites.pdf'])

%% Plot histogram of edits
fig = figure(33);

% Add a unicode character arrow between the edited bases
biochangechar = char(biochange);
firstletter = biochangechar(:,1);
arrow = repmat(char(0x2192),size(firstletter));
secondletter = biochangechar(:,2);
biochange2 = string([firstletter arrow secondletter]);
histogram(categorical(biochange2));


% Plot properties
fig.Units = "inches";
fig.Position = [0 0 2.14 1.67];
fontsize(fig,myfontsize,"Points")
fontname(fig,myfont);
%title('Overall biological editing changes')
ylabel('Number of occurences')
set(gca,'box','off')
exportgraphics(figure(33),[path.main '\Results\Overall Biological Editing Changes.pdf'])

%% Number of edits/cell vs mean editing percentage
fig = figure(34);

% Go through data_edits and find avg for each cell
u = unique(data_edits.CellID);

celltype = strings(size(u,1),1);
meaneditrate = zeros(size(u,1),1);
editspercell = zeros(size(u,1),1);
for i = 1:size(u,1)

    % Find indeces where this cell has edits listed in data_edits
    idx = ismember(data_edits.CellID,u(i));

    % Record this cell's celltype (Ib or Is)
    celltype(i) = data_edits{find(idx,1),"CellType"};

    % Calculate this cell's mean editing percentage
    meaneditrate(i) = mean(data_edits{idx,"FractionOfReadsEdited"});

    % Count how many edits were in this cell
    editspercell(i) = sum(idx);
end

% Delete outliers?
%outliers = find(isoutlier(meaneditrate,"quartiles"));
%meaneditrate = meaneditrate([1:98 100:end]);
%editspercell = editspercell([1:98 100:end]);
%celltype = celltype([1:98 100:end]);

% Find indeces for Ib and Is
idx_Ib = strcmp(celltype,"Ib");
idx_Is = strcmp(celltype,"Is");

% Plot
scatter(editspercell(idx_Ib),meaneditrate(idx_Ib),markersize,'.','CData',color_Ib)
R_Ib = corrcoef(editspercell(idx_Ib),meaneditrate(idx_Ib));
hold on
scatter(editspercell(idx_Is),meaneditrate(idx_Is),markersize,'.','CData',color_Is)
R_Is = corrcoef(editspercell(idx_Is),meaneditrate(idx_Is));
hold off


%title('Do cells with fewer editing sites have higher editing rate?')
ylabel(["Mean fraction of reads edited / cell" "(Each dot is one cell)"])
xlabel("Number of edits / cell")

% Lease-squares fit
h = lsline;
set(h(1),'color',color_Ib)
set(h(2),'color',color_Is)

%l = lsline;
%bestfitline = polyfit(editspercell,meaneditrate,1);
%bestfitlineval = polyval(bestfitline,min(editspercell):max(editspercell));
%hold on
%bestfitlineline = plot(min(editspercell):max(editspercell),bestfitlineval,'color','black','linewidth',1);
%hold off
L = legend("Ib","Is");
L.ItemTokenSize(1) = 10;
%[Rsq,Pvalfit] = corrcoef(editspercell,meaneditrate);

fig.Position = [0 1 2.7 2.3];
fontsize(fig,myfontsize,"Points")
fontname(fig,myfont);
exportgraphics(figure(34),[path.graphs '\EditSiteNum vs EditRate.pdf'])

%% Coverage and how many genes are expressed highly enough that they pass our thresholds?
% path.tpms = [path.main '\TPM data\'];
% tpmdata = readtable([path.tpms 'Jetti et al - Supp Table 1.xlsx'],'Range','A5:JC15032','Sheet','1b--Counts');
% tpmdata = [tpmdata(:,12:116) tpmdata(:,118:218)];
% 
% % For each gene, pass or fail if it has 10 genes with 10 reads minimum
% pass = zeros(size(tpmdata,1),1);
% Ib_nnz(i) = zeros(size(tpmdata,1),1);
% Is_nnz(i) = zeros(size(tpmdata,1),1);
% for i = 1:size(tpmdata,1)
% 
%     Ib = 1:105;
%     Is = 106:206;
% 
%     Ib_nnz(i) = nnz(tpmdata{i,Ib}>10);
%     Is_nnz(i) = nnz(tpmdata{i,Is}>10);
% 
%     if mod(i,100)==0
%     disp(i)
%     end
%     
% end
% 
% % Count how many genes had at least 10 cells in Ib or Is
% sum(Ib_nnz>10 | Is_nnz>10)


%% Read depth at cells without edits
fig = figure(35);

x = [data_summary{:,'Mean reads in Ib cells with edit'};...
    data_summary{:,'Mean reads in Is cells with edit'}];
y = [data_summary{:,'Mean reads in Ib cells without edit'};...
    data_summary{:,'Mean reads in Is cells without edit'}];

mdl = fitlm(x,y,'RobustOpts','andrews');
plot(mdl)

%scatter(x,y,markersize,'.');
title('Mean reads per edit')
xlabel('Mean reads in cells with edit')
ylabel('Mean reads in cells without edit')
xlim([0 150])
ylim([0 150])
axis square
legend off

hold on
refline(1,0)
hold off

exportgraphics(figure(35),[path.graphs '\Reads in cells without edit.pdf'])
%% Does difference in cells depend on reads
fig = figure(36);

yIb = data_summary{:,'Mean reads in Ib cells without edit'}-...
      data_summary{:,'Mean reads in Ib cells with edit'};
yIs = data_summary{:,'Mean reads in Is cells without edit'}-...
      data_summary{:,'Mean reads in Is cells with edit'};
xIb = data_summary{:,'Num of Ib cells without edit'}-...
      data_summary{:,'Num of Ib cells with edit'};
xIs = data_summary{:,'Num of Is cells without edit'}-...
      data_summary{:,'Num of Is cells with Edit'};

x = [xIb;xIs];
y = [yIb;yIs];

scatter(x,y,markersize,'.');
title('Does edit detection depend on read count?')
ylabel(["Difference in read counts","(Reads in cells without edit - reads in cells with edits)"])
xlabel(["Difference in cell count","(Num of cells without edit - cells with edit)"])
%xlim([0 150])
ylim([-500 500])
%axis square
legend off

exportgraphics(figure(36),[path.graphs '\Does edit detection depend on read count.pdf'])

%% MUSCLES VERSUS NEURONS

%% venn diagram of muscle and neuron edits
fig = figure(37);

% Run this block to find all instance where we detected at least 1 muscle
% edit
%index_Ib = data_summary.("Mean Ib editing") > 0;
%index_Is = data_summary.("Mean Is editing") > 0;
%index_M = data_summary.("Mean M editing") > 0;

% Run this block to find all instances where we dtected at least 10 muscle
% with >0.1 editing rate
% edits
index_Ib = data_summary.("Mean Ib editing") > 0 & data_summary.("Num of Ib cells with edit") > 10;
index_Is = data_summary.("Mean Is editing") > 0 & data_summary.("Num of Is cells with Edit") > 10;
index_M = data_summary.("Mean M editing") > 0.1 & data_summary.("Num of M samples with edit") > 10;

% Set up variables
y = false(size(data_summary,1),1);
x = [];

% Only Ib
y(:,1) = index_Ib & ~index_Is & ~index_M;

%Only Is
y(:,2) = index_Is & ~index_Ib & ~index_M;

%Only M
y(:,3) = index_M & ~index_Ib & ~index_Is;


%Ib+Is
y(:,4) = index_Ib & index_Is & ~index_M;

%Ib+M
y(:,5) = index_Ib & ~index_Is & index_M;

%Is+M
y(:,6) = ~index_Ib & index_Is & index_M;


%Ib+Is+M
y(:,7) = index_Ib & index_Is & index_M;


% Create category names for bar plot
catnames = ["Total" ...
            "Ib only" ...
            "Is only" ...
            "M only" ...
            "Ib+Is" ...
            "Ib+M" ...
            "Is+M" ...
            "Ib+Is+M"];

x = categorical(catnames);
x = reordercats(x,catnames);

% Find sum of each column to represenet total count per category
sums = [sum(y,'all') sum(y)];
%sums = [sums sums(1)-sum(sums(2:end))];

% Plot bar graph
b = bar(x,sums);

xlabel("Cell type")
ylabel("# of edits")
xtips1 = b(1).XEndPoints;
ytips1 = b(1).YEndPoints;
labels1 = string(b(1).YData);
text(xtips1,ytips1,labels1,'HorizontalAlignment','center',...
    'VerticalAlignment','bottom')


fig.Position = [0 1 2.7 2.3];
fontsize(fig,myfontsize,"Points")
fontname(fig,myfont);
exportgraphics(figure(37),[path.graphs '\Muscle--Num of edits.pdf'])

clear x y sums labels1 xtips1 ytips1 b

%% Save table of muscle edits
writetable(data_summary(index_M,:),...
    [path.tables '\Supplemental Table 7 - Muscle edits.xlsx'],...
    'WriteMode','overwrite','Sheet','All M edits')

writetable(data_summary(index_M & ~index_Ib & ~index_Is,:),...
    [path.tables '\Supplemental Table 7 - Muscle edits.xlsx'],...
    'WriteMode','overwrite','Sheet','M-only edits')

%% Boxcharts of top M edits
fig = figure(38);

% Create new table with just the edits to plot
index = data_summary.("Cell types with edit") == "NM";
tbl = data_summary(index,:);
tbl.('Mean N editing') = mean([tbl.("Mean Ib editing"),tbl.("Mean Is editing")],2);
tbl = sortrows(tbl,"Mean N editing",'descend');

% Create unique names for x labels
data = tbl.GenePart;
data = replace(data,"FivePrimeUTR","5'UTR");
data = replace(data,"ThreePrimeUTR","3'UTR");
tbl.UniqueName = append(tbl.("Edit Position"),"--",data,"--",tbl.Gene);
x = categorical(tbl.UniqueName);
x = reordercats(x,tbl.UniqueName);

% Plot
y = [tbl.('Mean Ib editing'),tbl.('Mean Is editing'),tbl.('Mean M editing')];
s = scatter(x,y,markersize,'.');

% Adjust plot appearance
s(1).CData = color_Ib;
s(2).CData = color_Is;
s(3).CData = [229 38 37]./255;
legend('Ib','Is','M','Location','northeast')
%xlabel('Gene')
ylabel('Mean fraction of reads edited')
xaxis=get(gca,'XAxis');
xaxis.FontSize = myfontsize;
xtickangle(45);


index_starIb = tbl.("P-val of Ib vs M editing") < 0.05;
starIb = strings(size(x,1),1);
starIb(index_starIb) = "*";
text(x,repmat(101,size(x,1),1),starIb,'HorizontalAlignment','center',...
    'VerticalAlignment','bottom','Color','blue')

index_starIs = tbl.("P-val of Is vs M editing") < 0.05;
starIs = strings(size(x,1),1);
starIs(index_starIs) = "*";
text(x,repmat(105,size(x,1),1),starIs,'HorizontalAlignment','center',...
    'VerticalAlignment','bottom','Color','red')

fig.Position = [0 1 7 3.8];
fontsize(fig,myfontsize,"Points")
fontname(fig,myfont);
ylim([0 1]);

exportgraphics(figure(38),[path.graphs '\Muscle--Editing rate.pdf'])

%% Calculate t test between neuron and muscle adar expression

if canonical == true
    % Extract ADAR transcripts per million for each single sequenced cell
    %path.tpms = 'C:\Users\Littleton Lab\Documents\Andres C drive\RNA editing\RNA editing code (compares RNAseq to DNAseq of parent flies)';
    tpmdata_adar = readmatrix('../TPM data/Jetti et al - Supp Table 1.xlsx','Sheet','1a--TPMs','Range','A1822:JC1822');
    tpmdata_adar_N = tpmdata_adar([12:116 118:218]);
    tpmdata_adar_M = tpmdata_adar([220:236 238:end]);
    
    [h,p] = ttest2(tpmdata_adar_M,tpmdata_adar_N);
end
%% Waterfall plots of top genes
fig = figure(39);

%cpx = FBgn0041605
% Load read depths (previously calculated in GATK for these locations)-----
if ~exist("readdepth","var")
    opts = delimitedTextImportOptions("NumVariables", 4);
    opts.DataLines = [2, Inf];
    opts.Delimiter = ["\t", " ", ":"];
    opts.VariableNames = ["Edit", "Position", "Cell", "Depth"];
    opts.VariableTypes = ["string", "double", "string", "double"];
    opts.ExtraColumnsRule = "ignore";
    opts.EmptyLineRule = "read";
    opts.ConsecutiveDelimitersRule = "join";
    opts = setvaropts(opts, ["Edit", "Cell"], "WhitespaceRule", "preserve");
    opts = setvaropts(opts, ["Edit", "Cell"], "EmptyFieldRule", "auto");
    
    filedir = "C:\Users\Littleton Lab\Documents\Andres C drive\RNA editing\GATK analysis - Muscle vs neurons\GATK output\Coverage\Read depth at specific locations\";
    batch1 = readtable(append(filedir,"neuron_RNAseq061819_coverage_at_specific_locations.txt"), opts);
    batch2 = readtable(append(filedir,"neuron_RNAseq082119_coverage_at_specific_locations.txt"), opts);
    batch3 = readtable(append(filedir, "neuron_RNAseq091219_coverage_at_specific_locations.txt"), opts);
    muscle = readtable(append(filedir, "muscle_coverage_at_specific_locations.txt"), opts);
    
    clear opts
    
    % Combine read depths into single table
    readdepth = [batch1; batch2; batch3; muscle];
    clear batch1 batch2 batch3 muscle
    
    % Change cellID to standard format
    readdepth.Cell = append("d",extractAfter(readdepth.Cell, "D19-"));
    
    % Combine Chr and Location
    readdepth.Location = append(readdepth.Edit,":",string(readdepth.Position));
end


% Load mastertable if it's not loaded yet----------------------------------
    if ~exist("masterTableFiltered","var")
    load([path.main 'GATK output\filtered edits.mat'])
    
    % Only save edits that passed filters 1 and 2 (min read depth and min qual)
    masterTableFiltered = masterTable(masterTable.Filter>2,{'CellID' 'CellType' 'Edit Position' 'FractionOfReadsEdited'});
    end

% Find IDs of cells that have edits, and sort so Ibs and Iss are together
cellnames = unique(string([data_edits.CellID data_edits.CellType]),'rows');
cellnames = sortrows(cellnames,2);
cellnames = num2cell(cellnames);

% For each cell, extract the fraction of reads edited at each site
numCells = size(cellnames,1);
for row = 1:numCells
    thisCellsEditsIndeces = strcmp(data_edits.CellID,cellnames{row});
    cellnames(row,3) = {data_edits.FractionOfReadsEdited(thisCellsEditsIndeces)};
end

% Find the average fraction of reads edited, and sort by this within Ib and
% Is
cellnames(:,4) = num2cell(cellfun(@mean,cellnames(:,3)));

indeces_Ib = strcmp([cellnames{:,2}],"Ib");
indeces_Is = strcmp([cellnames{:,2}],"Is");
indeces_M = strcmp([cellnames{:,2}],"M");

cellnames(indeces_Ib,:) = sortrows(cellnames(indeces_Ib,:),4);
cellnames(indeces_Is,:) = sortrows(cellnames(indeces_Is,:),4);

plotmuscles = false;
if plotmuscles == true
    cellnames(indeces_M,:) = sortrows(cellnames(indeces_M,:),4);
else
    cellnames(indeces_M,:) = [];
end

% cpx sites
titletext = 'Cpx sites';
genes = ["cpx";"cpx";"cpx"];
sites = ["3R:4297504";"3R:4297517";"3R:4297518"];

% highly edited N sites
% titletext = 'nAchRalpha5, RIM, slo, cac';
% genes = ["nAChRalpha5";"Rim";"slo";"cac"];
% sites = ["2L:14083516";"3R:17881419";"3R:24707423";"X:11977538"];

% highly edited M sites
% titletext = 'M editing sites - LSm3,Tim9b,Pstk,Cdc50';
% genes = ["LSm3";"Tim9b";"Pstk";"Cdc50"];
% sites = ["3R:23996648";"X:19618035";"X:19618035";"X:16441177"];

% M and N sites
% titletext = 'M and N sites';
% genes = ["CSN4";"CG15528";"Ald1"];
% sites = ["2R:7973697";"3R:30052032";"3R:26254381"];


% For each edit site to graph, add a col to cellnames:
% col 1: a list of editing rates for each cell (in current cellnames order)
% extracted from masterTableFiltered
for site = 1:size(sites,1)
    currentSite = sites(site);
    
    % Create temp var to hold each cell's edit rate for current site
    temp = nan(size(cellnames,1),1);


    % For each cell in cellnames, find the edit rate in masterTableFiltered
    for i = 1:size(cellnames,1)
        currentCell = cellnames{i,1};
    
            % Using logical indexing, search for a row in masterTableFiltered
            % with the current cellID and edit position
            index_ID = masterTableFiltered{:,'CellID'} == currentCell;
            index_ed = masterTableFiltered{:,'Edit Position'} == currentSite;
            index_ma = index_ID & index_ed;
        
            % Using the logical index, extract the edit rate
                % If there is a match, then we know there was an edit detected
                % with at least 10 reads. Record the editing rate.
                if sum(index_ma) == 1
                    temp(i,1) = 100*masterTableFiltered{index_ma,'FractionOfReadsEdited'};
    
                % If there is no match, then we don't know if it's bc there was
                % actually 0 editing in this cell in this location, or bc there
                % were no reads at this site. To solve this, query the list of
                % read depths at specific locations. If there are more than 10
                % reads, then the editing rate is 0. If there are less than 10
                % reads then there is not enough read depth to determine, so
                % leave nan.
                elseif sum(index_ma) == 0
    
                    index_ID2 = readdepth{:,'Cell'} == currentCell;
                    index_ed2 = readdepth{:,'Location'} == currentSite;
                    index_ma2 = index_ID2 & index_ed2;
    
                    if readdepth{index_ma2,'Depth'} > 10
                        temp(i,1) = 0;
                    end
                end
    end
    % Save the matrix into a cell in cellnames
    cellnames(:,site+4) = num2cell(temp);
end


% Sort cellnames by the first gene
cellnames(indeces_Ib,:) = sortrows(cellnames(indeces_Ib,:), 5,'ascend');
cellnames(indeces_Is,:) = sortrows(cellnames(indeces_Is,:), 5,'ascend');

if plotmuscles == true
    cellnames(indeces_M,:) = sortrows(cellnames(indeces_M,:), 5,'ascend');
end

% Create layout
t = tiledlayout(size(sites,1),1);
H = gobjects(size(sites,1),1);

for i = 1:size(sites,1)
    H(i)=nexttile;

    % Make a stacked bar plot
    plotData = cell2mat(cellnames(:,i+4));
    plotData(:,2) = 100-plotData;
    plotData(:,3) = 0;
    b = bar(plotData,1,'stacked');
    b(3).FaceColor = 'white';
    b(3).EdgeColor = 'black';

    xticks([105 206])
    
    title(sites(i))
    xticklabels([])
    set(gca,'TickDir','out')

    if i == size(sites,1)

        if plotmuscles == true
            xticklabels({'<--Ib Is-->','<--Is M-->'})
        else
            xticklabels({'<--Ib Is-->',''})
        end
        %plot(nan, nan, 'DisplayName', "mean m/s");
        legend('Edited Reads','Unedited Reads','Cell with <10 reads','Location','southoutside')
    end

end

fig.Position = [0 1 3 4];
%title(t,titletext)
ylabel(t,'Percent of reads')
%xlabel(t,'Individual cells/muscle samples')
fontsize(fig,myfontsize,"Points")
fontname(fig,myfont);
exportgraphics(figure(39),[path.graphs '\EditingRateSpecificSitesAcrossCells-' titletext '.pdf'])

%% Split edits into N or M categories
g = groupsummary(data_summary,"Cell types with edit");

indexN = contains(data_summary.("Cell types with edit"),"N");
groupsummary(data_summary(indexN,:),"BiologicalBaseChange")

groupsummary(data_summary(indexN,:),"GenePart")

%% Back to back horizontal bar graph comparing TPM expression to editing

% Import data
data = readtable([path.tables '\Supplemental Table 6 - Differences between Ib and Is.xlsx']);
data = sortrows(data,'MeanIbEditing_MeanIsEditing');
data = data([1:3 (end-3):end],:);
numentries = size(data,1);

% Axes 1 - Plot TPMs
fig = figure(40);

ax_front = axes;
x = 1:1:numentries;
y = [data.GeneTPMInIb data.GeneTPMInIs];
b = barh(y);
b(1).FaceColor = color_Ib;
b(2).FaceColor = color_Is;
axmax = max(xlim);
axis([-axmax axmax 0.5 numentries+0.5])
%ax_front.XTick = -axmax:1:axmax;
%ax_front.XTickLabel = [repmat-axmax:1:axmax;
L1 = legend('Ib','Is');
L1.Position = [-axmax numentries 1 1];
L1.AutoUpdate = 'off';
axis off


% Axes 2 - Plot RNA editing levels
hold on
ax_back = axes;
bb = barh([-data.MeanIbEditing -data.MeanIsEditing]);
hold off
bb(1).FaceColor = color_Ib;
bb(2).FaceColor = color_Is;
axis([-1 1 0.5 numentries+0.5]);

set(gca, 'xtick', [-1:0.2:0,1/axmax:1/axmax:1]);
set(gca, 'xticklabel', [1:-0.2:0,1:1:axmax])

axes(ax_front)


% Add Gene name labels
x = repmat(0,numentries,1);
offset = 0.4;
y = 1+offset:1:size(data,1)+offset;
text(x,y,string(data.Gene))

% Add significance stars
sig_tpm = repmat('ns',numentries,1);
x = repmat(axmax,numentries,1);
text(x,y-offset,sig_tpm);

sig_edit = data.P_valOfIbVsIsEditing;
sig_edit_label = repmat("*",numentries,1);
sig_edit_label(sig_edit>0.05) = "*";
sig_edit_label(sig_edit>0.005) = "**";
sig_edit_label(sig_edit>0.0005) = "***";
sig_edit_label(sig_edit>0.00005) = "****";
t = text(-x,y-offset,sig_edit_label);
set(t,'HorizontalAlignment','center');
set(t,'Rotation',90);
%t(:).Rotation = deal(90);
%repmat(90,numentries,1);

% Add x axis labels
label(1) = text(axmax / 2, -0.3, 'TPMs','HorizontalAlignment', 'center');
label(2) = text(-axmax/ 2, -0.3, 'Mean RNA editing','HorizontalAlignment', 'center');

% Figure formatting
fig.Position = [0 1 3 4];
ax_back.YAxis.Visible = 'off';
ax_back.Box = 'off';
fontsize(fig,myfontsize,"Points")
fontname(fig,myfont);

exportgraphics(figure(40),[path.graphs '\Genes with similar TPM exp but diff editing bar chart.pdf'])

%% Edit sites with >0.9 editing in Ib or Is

% Include sites with editing rates above this threshold in Ib OR Is
threshold = 0.9;

% Create table of just the values that passed the threshold
index1 = (data_summary.("Mean Ib editing") > threshold) | ...
        (data_summary.("Mean Is editing") > threshold);
index2 = data_summary.MutationType == "Missense";
high = data_summary(index1&index2,:);

% Calculate the number of unique sites and unique genes
disp(["There are " num2str(size(high,1)) " editing sites with rates over " num2str(threshold) ...
    ", consisting of " num2str(size(unique(high.Gene),1)) " unique genes."])
groupsummary(high,'PreviouslyKnownRNAeditingSite')
writetable(high,[path.tables '\Supplemental Table 5 - Highest edited sites leading to amino acid substitution.xlsx'],'WriteMode','overwrite')
%% Functions
function fig = boxswarmchart(fig,sorted,data_edits,numEditsToPlot,color_Ib,color_Is,markersize)

% If numEditsToPlot is larger than the table of sorted edits to plot,
% change the numEditsToPlot to be equal to the table size
if size(sorted,1)<numEditsToPlot
    numEditsToPlot = size(sorted,1);
end


points = table();

% Go through the top edits and record individual edit fractions
for i = 1:numEditsToPlot
    index_Ib = (strcmp(data_edits.FlybaseID, sorted{i,"FlybaseID"})) & ...
            strcmp(data_edits.("Edit Position"),sorted{i,"Edit Position"}) & ...
            (strcmp(data_edits.CellType, 'Ib'));
    index_Is = (strcmp(data_edits.FlybaseID, sorted{i,"FlybaseID"})) & ...
        strcmp(data_edits.("Edit Position"),sorted{i,"Edit Position"}) & ...
        (strcmp(data_edits.CellType, 'Is'));

    newdata = data_edits(index_Ib,[ "CellType" "Gene" "Edit Position" "FractionOfReadsEdited"]);
    points = [points; newdata];

    newdata = data_edits(index_Is,[ "CellType" "Gene" "Edit Position" "FractionOfReadsEdited"]);
    points = [points; newdata];
end

% Combine Gene and position into a naming variable for plotting
%points.Name = append(string(points.Gene),"(",string(num2str(points.("Edit Position"))),")");
points.Name = append(string(points.Gene),"(",points.("Edit Position"),")");
nameorder = append(string(sorted{1:numEditsToPlot,"Gene"}),"(",sorted{1:numEditsToPlot,"Edit Position"},")");

% Plot
figure(fig);
fig.Position = [531 918 1038 420];
index = strcmp(points.CellType,'Ib');
x_Ib = categorical(points.Name(index),nameorder);
y_Ib = points.FractionOfReadsEdited(index);

b = boxchart(x_Ib,y_Ib);
b.BoxFaceColor = color_Ib;
b.WhiskerLineColor = color_Ib;
b.MarkerStyle = '.';

hold on
index = strcmp(points.CellType,'Is');
x_Is = categorical(points.Name(index),nameorder);
y_Is = points.FractionOfReadsEdited(index);

b = boxchart(x_Is,y_Is);
b.BoxWidth = 0.2;
b.BoxFaceColor = color_Is;
b.WhiskerLineColor = color_Is;
b.MarkerStyle = '.';


s = swarmchart(x_Ib,y_Ib,'.');
s.SizeData = markersize;
s.MarkerEdgeColor = color_Ib;

s = swarmchart(x_Is,y_Is,'.');
s.SizeData = markersize;
s.MarkerEdgeColor = color_Is;

f=get(gca,'Children');
legend(f(3:4),'Is','Ib','Location','northeastoutside')
hold off

ylabel('Fraction of reads edited')
xlabel('Gene and position of edit')


end
```
